# Multiplexed neuropeptide mapping in ant brains integrating microtomography and 3D mass spectrometry imaging

**DOI:** 10.1101/2022.11.02.514707

**Authors:** Benedikt Geier, Esther Gil-Mansilla, Zita Liutkeviciute, Roland Hellinger, Jozef Vanden Broeck, Janina Oetjen, Manuel Liebeke, Christian W. Gruber

## Abstract

Neuropeptides are important regulators of animal physiology and behavior. Hitherto large-scale localization of neuropeptides mainly relied on immunohistochemical methods requiring the availability of antibody panels, while another limiting factor has been the brain’s opacity for subsequent light or fluorescence microscopy. To address these limitations, we integrated high-resolution mass spectrometry imaging (MSI) with microtomography for a multiplexed mapping of neuropeptides in two evolutionary distant ant species, *Atta sexdens* and *Lasius niger*. For analyzing the spatial distribution of chemically diverse peptide molecules across the brain in each species, the acquisition of serial mass spectrometry images was essential. As a result, we have comparatively mapped the 3D distributions of eight conserved neuropeptides throughout the brain micro-anatomy. We demonstrate that integrating the 3D MSI data into high-resolution anatomy models can be critical for studying organs with high plasticity such as brains of social insects. Several peptides, like the tachykinin-related peptides TK1 and TK4, were widely distributed in many brain areas of both ant species, whereas others, for instance myosuppressin was restricted to specific regions only. Also, we detected differences at the species level; many peptides were identified in the optic lobe of *L. niger*, but only one peptide (ITG-like) was found in this region in *A. sexdens*. Our approach provides the basis for investigating fundamental neurobiological processes by visualizing the unbiased 3D neurochemistry in its complex anatomic environment.

**Significance statement:** Until recently, the inability to detect entire molecules such as neuropeptides within their spatial biological context and simultaneously link their occurrence to anatomically and physiologically relevant areas has limited our understanding of complex neurochemical processes. This situation has now changed dramatically with the optimization of a new multiplexed imaging method based on mass spectrometry, which enables us to study previously invisible processes at the microscopic scale. With the marriage of mass spectrometry imaging and microtomography, we show that it has become possible to build high-resolution maps of neuropeptides in complex anatomical structures as small as ant brains. These maps, embedded in the 3D neuroanatomy, expand the understanding of the spatial organization of brain chemistry and provide a baseline for neurobiological and neurochemical studies.

## Introduction

Neuropeptides are small molecules that act as extracellular signals regulating important biological processes, such as metabolism, development, reproduction, behavior and learning. The origin of neuropeptides is deeply rooted in metazoan evolution and likely emerged at very early stages of nervous system development (1).

In the animal kingdom, insects appeared around 479 million years ago (2) and represent the most diverse group in terms of numbers of described species. This taxon has an enormous significance with respect to ecology, biology and economy. In fact, insects participate in ecosystem maintenance, act as vectors of diseases, parasitize other animals and plants, pollinate our fruits or eat our crops with drastic impacts on agriculture. Amongst insects, social insects such as ants play a vital role in ecosystem functioning worldwide. Ants in particular display a distinguished social colony organization between species and unique caste systems within species, making them an emerging model system that is highly suitable for studying neuropeptide signaling (3). For instance, individuals from the same species belonging to differently sized colonies can exhibit a different brain anatomy depending on the tasks they develop (4, 5). To understand the molecular processes underlying the social behavior in different developmental stages, castes and species, a link has to be established between neuropeptide chemistry and brain anatomy at the level of individual organisms.

Recently, mining of published genome and transcriptome sequences of ants and other insects has resulted in the prediction of hitherto undescribed neuropeptides (6, 7). Their role, tissue expression and distribution remain unknown, except for a few, evolutionarily and behaviorally distinct model species (6, 8). Elucidating the biological function of neuropeptides could provide targets for insect pest management (9) and drug development (10, 11).

Detection and localization of neuropeptides is classically achieved by immunohistochemistry and *in situ* hybridization (12-15). However, these methods require the production of specific labels such as antibodies or complementary genetic probes, respectively. Particularly for multi labeling approaches (*e.g*., for detection of a set of neuropeptides), design and testing of the individual labels is highly time-consuming and conventionally limited by the number of fluorophores within a single labeling experiment.

Mass-spectrometry imaging (MSI) in contrast is a multiplexed and label-free technique, which can map the spatial distribution and relative abundance of hundreds to thousands of molecules from the same tissue section (16). Over the last decade, matrix-assisted laser desorption/ionization (MALDI)-MSI has revolutionized spatial biology (17). The field of neurochemistry has tremendously profited from the ability to detect a variety of molecular species, such as lipids and neurotransmitters, within a single measurement (18). The localization of neuropeptides with MALDI-MSI in insects is an emerging application and includes studies of the Jonah crab (19, 20), the desert locust (21) and two social insects: the African honeybee (*Apis melifera*) (22) and the desert ant (*Cataglyphis nodus*) (23).

One major challenge using MSI on sub-millimeter scale and complex tissue samples like insect brains is the translation of the generated ion maps into histology-connected chemical maps. Non-invasive 3-dimensional (3D) imaging techniques such as magnetic resonance tomography (MRT) in combination with MSI provided an improved integration of molecular and anatomic visualization data (24-26). 3D models have been used as virtual atlases, providing an anatomic framework for the co-registration with datasets of other imaging techniques, such as MS- and fluorescence imaging data (27).

For resolving the neuroanatomy of millimeter-sized arthropods, MRT does not provide the required spatial resolution. Therefore, microscopy-based imaging, such as confocal- and/or laser scanning microscopy has been widely used for insect neuroimaging (28). However, the penetration depth of the microscopy laser, for instance is limited to 100-300 μm in clear tissue samples and even less in opaque specimens, which means that penetrating the different types of head capsules that encase the arthropods’ brains, if not dissected, poses a substantial challenge.

Micro-computed tomography (μCT) is a X-ray-based, non-invasive 3D imaging technique capable of penetrating through any biological specimen and, unlike most microscopy-based methods, μCT provides the same spatial resolution along x, y and z axes (29, 30). In insects, other arthropods and even in fossils high-resolution 3D μCT imaging has been used to resolve taxonomic relations and to study the functional morphology (31-34). Although μCT is X-ray based and not limited in penetration depth, X-rays are poorly attenuated through animal soft tissues. Similar to electron microscopy, tissues are conventionally contrasted with heavy element agents (35). The development of more elaborate contrasting protocols (36, 37) and advances in spatial resolution below 1 μm have promoted applications of μCT beyond small-animal imaging, for instance with micro-scale 3D neuroanatomy reconstruction for the honey bee (29, 38). Furthermore, faster tomographic reconstructions now allow for effective screenings of small animal models supporting physiological and developmental studies (30, 39-42).

Given the advances in MALDI-MSI and μCT, we reasoned that combining both techniques could provide a powerful link for complementing neuropeptide chemistry with brain anatomy to build maps that advance our understanding of neurophysiology across castes and species of ants, and potentially many other animals.

In this study, we optimized a spatial peptidomics procedure by combining MALDI-MSI and μCT in two ant species, the leafcutter ant *Atta sexdens* and the black garden ant *Lasius niger*, resulting in high-resolution molecular neuropeptide mapping and anatomical 3D imaging. This correlative approach enabled us to integrate the spatial distribution of neuropeptides from all planes of the ant brain into a 3D anatomy model and link subregions of the brain with their specific neuropeptide chemistry. We advance previous separate MSI and μCT datasets (27, 43) by integrating them in a full 3D approach that provides unprecedented detail of ant brain anatomy in correlation to neuropeptide distribution.

## Results and Discussion

### Development of a workflow for neuropeptide imaging in social insects

We developed a combined methodological approach to link 3D spatial peptidomics to full brain microanatomy (see **Fig. 1**). Thereby, we focused on two species from different positions in the ant evolutionary tree (44) which display substantial differences in head morphology and social behavior. The first step of our approach was the identification of neuropeptides by dereplication present in each of the two model ant species, *L. niger* and *A. sexdens*. We mined four available ant genome data sets to predict neuropeptide sequences of the two model species in this study. From the amino acid sequences of each neuropeptide, we calculated the theoretical *m/z* values. These target *m/z* values were subsequently mapped using MALDI-MSI data from on a series of consecutive ant brain sections. In parallel, a comparable head in terms of size and caste of each species was contrasted and imaged by μCT, resulting in a 3D anatomical model. Finally, the spatial peptidomics data was manually co-registered into the anatomic 3D model in a virtual space resulting in a 3D atlas correlating anatomy with neuropeptide biochemistry for direct comparison and localization.

**Figure 1.**
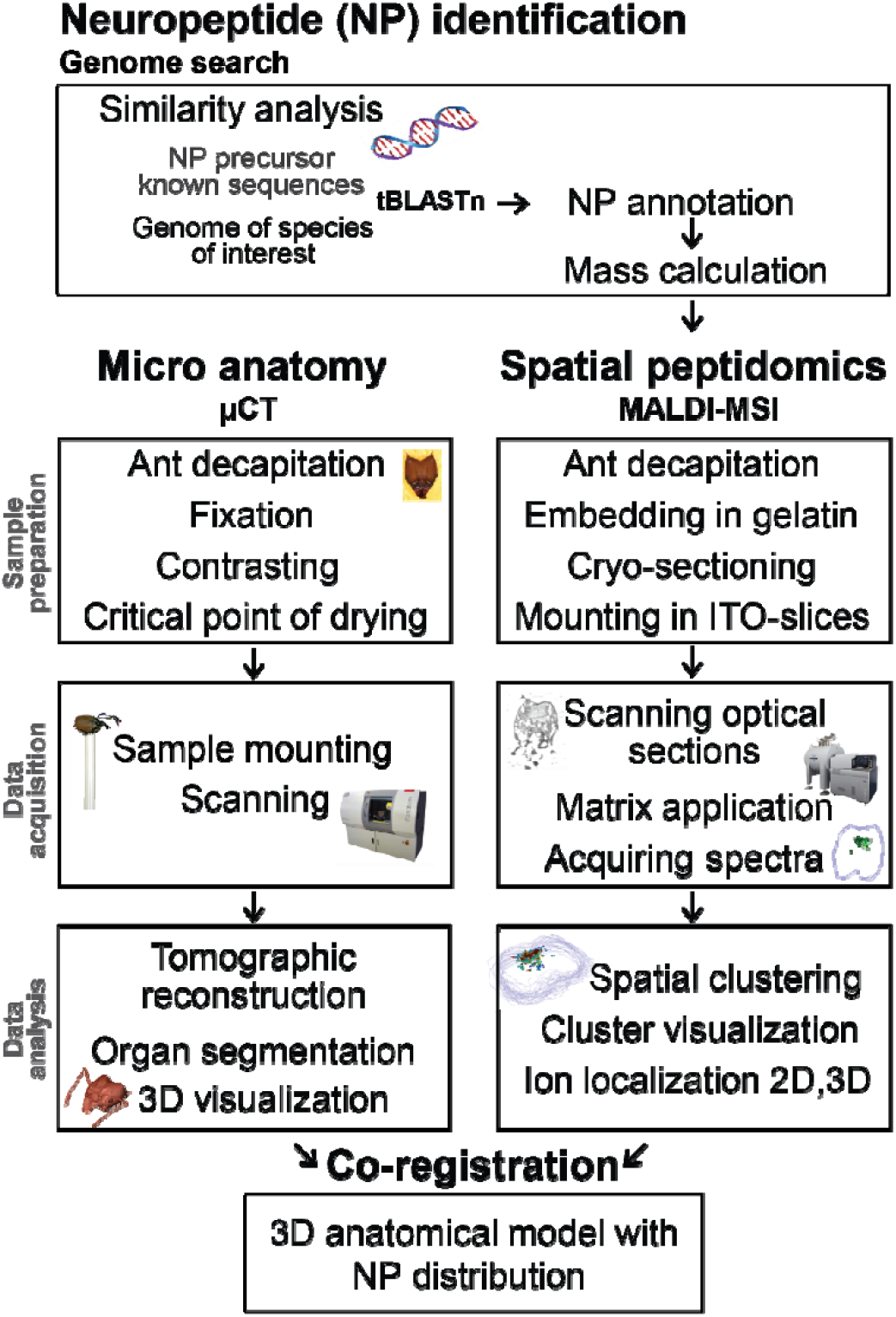
Workflow for genome-based neuropeptide identification and localization with correlative 3D imaging of brain anatomy and spatial peptidomics. Separate sample preparation, imaging and data analysis pipelines of the micro-computed tomography (μCT) and matrix-assisted laser desorption ionization mass spectrometry imaging (MALDI-MSI) data; micro-anatomy (via μCT) and spatial peptidomics (MALDI-MSI) were acquired from two separate samples of the same species and comparable life stages; co-registration of the μCT and MALDI-MSI datasets in AMIRA™ was based on manual expert annotation by matching brain areas between virtual μCT slices and optical images of the tissue sections used for MALDI-MSI.

### In silico prediction, sequencing and in situ detection of ant neuropeptides

For the *in silico* predictions, we compared published neuropeptide precursor sequences of *Acromyrmex echinatior* (45) and *Camponotus floridanus* (46) with the genome of *L. niger* (47) on one hand and *A. cephalotes* (48), a closed species to *A. sexdens*, on the other hand as the genome and transcriptome of *A. sexdens* ant is not available. In the genome of each species, we discovered 30 neuropeptide precursors, which encode 58 neuropeptides in *L. niger* and 57 in *A. cephalotes* (**Suppl. Table S1**). Furthermore, we confirmed the allatostatin A, myosuppressin, tachykinin-related peptide, and short neuropeptide F precursor sequences by molecular cloning to further validate the *in-silico* prediction of neuropeptide dataset (**Suppl. Table S1** and **Suppl. Information S1**). Particular *L. niger* neuropeptide sequences (13 of in total 15), which were encoded by these four cloned precursors, matched the ones obtained via *in silico* prediction. After comparing mature peptide sequences of different ants, we found that myrmicine species *A. echinatior* and *A. cephalotes* share 65% of the peptides with identical sequences, whereas *A. echinatior* and the formicine *L. niger* – evolutionary more distant species (49) – exhibited only 25% sequence identity at neuropeptide level. Since *A. cephalotes* and *A. sexdens* belong to the same genus we could expect high sequence conservation and neuropeptide similarities. After cloning four precursors and a comparative analysis of the neuropeptide sequences, we confirmed that the amino acid sequences for 10 out of 13 peptides were identical in both species, resulting overall in 76% neuropeptide identity between the two model ant species. We found the biggest difference within the precursor of tachykinin-related peptides in *A. sexdens*, which was 129 amino acids shorter compared to *A. cephalotes* and lacked two out of seven tachykinin-related peptides. Additionally, the tachykinin-related peptide TK7 in *A. sexdens* was not identical to TK7 in the other species but contained one additional serine (**Suppl. Fig. S1**). Our approach showed that high quality prediction of neuropeptides by *in silico* mining can be complemented through confirmation of peptide-specific masses via MALDI-MSI analysis.

### 3D Localization of neuropeptides in situ with serial MALDI-MSI (spatial peptidomics)

Neuropeptides are conventionally either detected in bulk brain extracts, using multiplexed approaches such as liquid chromatography-mass spectrometry (LC-MS/MS) or are visualized individually through specific fluorescent labels using immunohistochemistry (IHC). Using a MSI setup with high-mass resolving power, a magnetic resonance mass spectrometry (MRMS) instrument for untargeted MALDI-MSI enabled us to measure and compare the exact molecular mass of each detected neuropeptide in the tissues. MALDI-MSI enabled us to simultaneously resolve the spatial distribution of up to 15 neuropeptides from measurements of multiple sections derived from the two ant species *L. niger* and *A. sexdens*. Extending MALDI-MSI into a serial approach for spatial neuropeptidomics revealed the need for 3D MSI to cover the spatial diversity of neuropeptides throughout different regions of the brain. The lists of detected molecular masses were used to target the genome-predicted neuropeptide masses (**Suppl. Table S1)** of *L. niger* and *A. sexdens* and to annotate and localize each of the neuropeptides (see **Table 1**).

**Table 1.** Detected neuropeptides in the two analyzed ant species.

| *Lasius niger* |  |  |  |  | *Atta sexdens* |  |  |  |
| --- | --- | --- | --- | --- | --- | --- | --- | --- |
| Neuro-peptide | Peptide sequence<sup>†</sup> | HRM. [M+H]<sup>+</sup> |  |  | Peptide sequence | HRM. [M+H]<sup>+</sup> |  |  |
|  |  | Calc. | Mea-sured | Error [ppm] |  | Calc. | Mea-sured | Error [ppm] |
| *Allatostatin A (AST A)* |  |  |  |  |  |  |  |  |
| AST A1 | LPLYNFGI* | 935.535 | n.f. |  | LPLYTFGI*<sup>†</sup> | 922.540 | n.f. |  |
| AST A2 | TRPFSFGI*<sup>†</sup> | 923.510 | 923.513 | 3.2 | TRQFSFGI*<sup>†</sup> | 954.516 | 954.516 | 0 |
| AST A3 | LRDYRFGI*<sup>†</sup> | 1038.584 | 1038.584 | 0 | LRNYDFGI*<sup>†</sup> | 996.526 | n.f. |  |
| AST A4 | GGKPFSFGI* | 908.499 | n.f. |  | GNHQFGFGI*<sup>†</sup> | 975.480 | n.f. |  |
| AST A5 | GWKLATGETAVS* | 1218.648 | n.f. |  | VWKLATGETAVS*<sup>†</sup> | 1260.695 | n.f. |  |
| *CAPA* |  |  |  |  |  |  |  |  |
| CAPA1 | SAGLVYPRI* | 1071.631 | 1071.634 | 2.8 | SAGLVAYPRI* | 1045.615 | n.f. |  |
| CAPA2 | ALGIIHQPRI* | 1116.700 | 1116.704 | 3.6 | AFGIIHKPRIG* | 1207.742 | n.f. |  |
| *Corazonin (CRZ)* |  |  |  |  |  |  |  |  |
| CRZ | pQTFQYSRGWTN* | 1369.629 | n.f. |  | pQTFQYSRGWTN* | 1369.629 | 1369.631 | 1.4 |
| *Ecdysis triggering hormone (ETH)* |  |  |  |  |  |  |  |  |
| ETH1 | DEVPAFFLKIAKIPTLPRV* | 2153.284 | 2153.286 | 0.9 | EEVPAFFLKIAKIPTLPRV* | 2167.300 | n.f. |  |
| IDLSRF-like | IDLSRFYGHINT | 1435.733 | 1435.736 | 2.0 | IDLSRFYGHFNT | 1469.717 | 1469.720 | 2.0 |
| ITG-like | ITGQGNRLF | 1005.548 | 1005.551 | 3.0 | ITGQGNRLF | 1005.548 | 1005.548 | 0 |
| *Myosuppressin (MS)* |  |  |  |  |  |  |  |  |
| MS | pQDVDHVFLRF*<sup>†</sup> | 1257.638 | 1257.642 | 3.4 | pQDVDHVFLRF*<sup>†</sup> | 1257.638 | 1257.639 | 0.7 |
| *Neuropeptide-like precursor 1 (NPLP1)* |  |  |  |  |  |  |  |  |
| NPLP1 2 | NVGTLDARDFALPT* | 1373.754 | 1373.751 | 2.2 | NVGALARDFALPT* | 1343.743 | n.f. |  |
| NPLP1 3 | HIASVARDHGLPN* | 1385.740 | 1385.746 | 4.3 | HIGSVLRDYSTMS* | 1464.726 | n.f. |  |
| NPLP1 4 | NIGSLARQSTLPSN* | 1456.787 | 1456.794 | 4.5 | NIGSLARQSMLPIS* | 1485.821 | 1485.822 | 0.7 |
| NPLP1 6 | NVAALARDSSLPY* | 1375.733 | 1375.732 | 0.7 | NVAALARDSSLPY* | 1375.733 | 1375.735 | 1.5 |
| *Orcokinin (OK)* |  |  |  |  |  |  |  |  |
| OK3 | NFDEIDRVGWGGFV | 1610.760 | n.f. |  | NFDEIDRAGWGGFV | 1582.728 | 1582.732 | 2.5 |
| *Pyrokinin (PK)* |  |  |  |  |  |  |  |  |
| PK1 | TTAQEITSGMWFGPRL* | 1793.900 | n.f. |  | TTQDITSGMWFGPRL* | 1708.848 | 1708.852 | 2.3 |
| *Short neuropeptide F (sNPF)* |  |  |  |  |  |  |  |  |
| sNPF | SPSLRLRF*<sup>†</sup> | 974.589 | 974.592 | 3.1 | SPSLRLRF*<sup>†</sup> | 974.589 | 974.589 | 0 |
| *Tachykinin-related peptides (TK)* |  |  |  |  |  |  |  |  |
| TK1 | APMGFQGMR*<sup>†</sup> | 993.476 | 993.480 | 2.8 | APMGFQGMR*<sup>†</sup> | 993.476 | 993.476 | 0 |
| TK2 | n.f. |  |  |  | APKGFQGMR*<sup>†</sup> | 990.530 | 990.531 | 1.0 |
| TK3 | LLAMGFQGIR*<sup>†</sup> | 1104.635 | n.f. |  | ASMGFQGMR*<sup>†</sup> | 983.455 | 983.456 | 1.0 |
| TK4 | TVMGFQGMR*<sup>†</sup> | 1025.502 | 1025.504 | 2.0 | TLMGFQGMR*<sup>†</sup> | 1039.518 | 1039.518 | 0 |
| TK7 | AAMGFYGTR*<sup>†</sup> | 972.472 | 972.472 | 0 | AALGFYGTR*<sup>†</sup> | 954.516 | 954.516 | 0 |

<sup>†</sup> confirmed by cloning; \*amidated C-terminus; p – pyroglutylated N-terminus; HRM – high resolution mass; calc. – calculated; n.f. – not found

The annotated neuropeptides were encoded by 10 prepropeptide genes in the central nervous system. For *L. niger*, we annotated 16 neuropeptides by MALDI-MSI, which encompass 27.5% of the 58 neuropeptides predicted by genome mining. For 8 neuropeptide sequences derived from *L. niger*, we confirmed the expression by precursor cloning and sequencing. For *A. sexdens*, the searches detected 15 *m/z*-values in the MALDI-MSI data, which matched the predicted neuropeptide masses, of which 12 were validated by cloning and sequencing of the precursor (**Table 1**). A limitation arose due to the same atomic composition of C_44_H_68_N_13_O_11_ for allatostatin (Ast) A2 and tachykinin-related peptide (TK) 7 of *A. sexdens*. Therefore, we could not differentiate these two neuropeptides with MALDI-MSI based on detected high-resolution *m/z* values on the MS1 level in this ant species. In summary, 58 neuropeptides were predicted by genome mining of the related species, of which as a further layer of evidence we cloned and sequenced 12 neuropeptides in *A. sexdens*. From these 12 sequenced neuropeptides in *A. sexdens*, we annotated eight by exact mass determination (see **Table 1**).

Using spatial peptidomics of ant brains we were able to account for a total of 8 neuropeptides ‘shared’ between both species, namely IDLSRF-like peptide, ITG-like peptide, myosuppressin (MS), neuropeptide-like precursor 1 derived peptides (NPLP1) 4 and 6, short neuropeptide F (sNPF), tachykinin-related peptides (TK) 1 and 4 (see **Fig. 2A**). To maintain a comparable measurement time to ensure sample stability to compare both species, for *L. niger* we acquired spatially resolved mass spectra with a lateral resolution of 40-times 40 μm from 44 consecutive sections, and for *A. sexdens* data from 32 sections with a pixel size of 30 μm. Neuropeptide signals were not present in all measured sections, demonstrating that classical two-dimensional (2D) MALDI MSI is not sufficient to cover neuropeptide distribution, even in specimens with head sizes as small as one millimeter. The high mass resolution MRMS data enabled a clear assignment of *m/*z values to the neuropeptides by exact mass determination (**Table 1**) and is shown exemplarily for TK1 in *L. niger* (inset **Fig. 2B**, mass error 2.8 ppm) and ITG-like peptide in *A. sexdens* (inset **Fig. 2C**, mass error 0.5 ppm). This methodology illustrates the possibility to visualize hundreds of different molecules through the multiplexed MS analysis as reflected in the mean spectra of all sections for *L. niger* and *A. sexdens* (see **Fig. 2B** and **C**).

**Figure 2.**
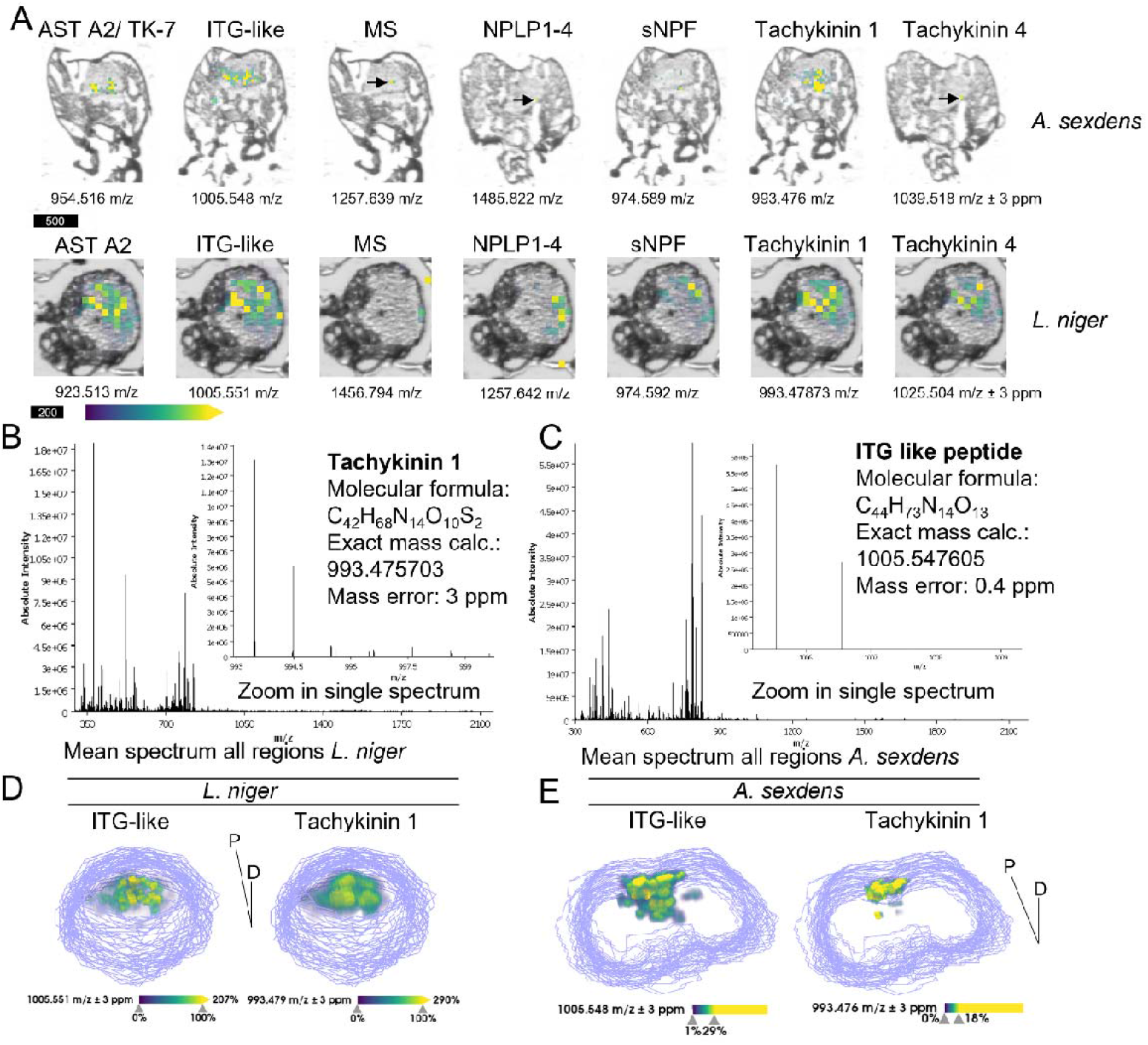
Mass-spectrometry imaging maps of neuropeptides in ant brain tissue sections. (**A**) Neuropeptide maps for seven different peptides detected in both, *A. sexdens* (upper panel, scale bar 500 μm) and *L. niger* (lower panel, scale bar 200 μm). Different single sections were selected for *A. sexdens* as neuropeptides occurred not in all measured sections. A representative central head section was selected for *L. niger*. For both species, neuropeptide signal intensity was visualized by false-color coding in overlay images with a low-resolution scan of the tissue sections using a viridis coloring scheme. Linear transparency was applied to make low intensity pixels transparent. Arrows indicate single pixels with neuropeptide signals being detected, all peptide ions were shown as [M+H]^+^ (see table 1 for *m/z* values) (**B**) Mean MRMS mass-spectrum of all measured regions as defined during the setup of the measurement for *L. niger*, insert shows tachykinin 1 ([M+H]^+^, calc. *m/z* 993.476, measured *m/z* 993.479, error 2.8 ppm). (**C**) Mean mass-spectrum for *A. sexdens*, inset shows ITG-like peptide ([M+H]^+^, calc. *m/z* 1005.5482, measured *m/z* 1005.5487, error 0.5 ppm). (**D, E**) Three-dimensional maps of ITG-like neuropeptide and tachykinin 1 in *L. niger* (**D**) and *A. sexdens* (**E**). Data shown here are from 44 central head sections of *L. niger* and 32 sections of *A. sexdens*. The dorso-ventrally cut sections were digitally stacked on each other by manual coregistration based on the low-resolution grayscale scans of the individual sections in SCiLS Lab 2023a. The section border is represented as gray line. Neuropeptide 3D-distributions were visualized in volume mode using a viridis color map. Low intensity pixels were made transparent. AST A2, allatostatin A2; MS, myosuppressin; NPLP1-4, neuropeptide like precursor 1-4; sNPF, short neuropeptide F; Tachykinin 1, Tachykinin 4, TK-7, tachykinin-related peptide 1,4 or 7. Orientation: P, posterior; D, dorsal.

We used spatial clustering of the full MSI data including all signals from peptides to lipids to group the molecular images after spatial and spectral similarities (**Suppl. Fig. S2**). The spatial clusters showed a clear separation between the local chemistry of tissues surrounding the brain (*i.e*., musculature) and the brain itself, as verified through overlays with optical scans of the same sections (**Suppl. Fig. S2**). All detected neuropeptides were centered in the brain-correlated spatial cluster indicating that each 3D MSI dataset contained the in-depth neuropeptidomic composition of the ants’ brains (see **Fig. 2D** and **2E** for ITG-like peptide and TK1 for *L. niger* and *A. sexdens*). However, without precise knowledge on the 3D orientation and co-localization between peptides and specific regions of the brain, analysis and interpretation of the different neuropeptide distributions remained challenging. To further exploit the results of the 3D MSI data, we wanted to allocate the 3D neuropeptide chemistry to well-studied (sub)regions of the ants’ brains, such as antennal- and optical lobes or mushroom bodies.

### Anatomical model of ant brains based on μCT data

Using non-invasive μCT we generated micrometer-scale models of two ant heads and their interior structures from developmental stages and castes of *L. niger* and *A. sexdens* similar to the samples used for spatial peptidomics. These high-resolution anatomical μCT models provided the 3D framework to precisely co-register the serial MSI data.

For μCT sample preparation we adapted a combination of protocols using phosphotungstic acid (36, 50) and added subsequent critical point drying (CPD) (38). CPD is conventionally used to preserve tissues of small arthropods for μCT scanning for an improved signal-to-noise (s/n) ratio as liquids from non-tissue filled compartments around the brain are removed. Phosphotungstic acid staining in combination with CPD enabled us to scan the intact heads of both ant species and resolve substructures of the ant brain down to a sub-micron level (see **Fig. 3**).

**Figure 3.**
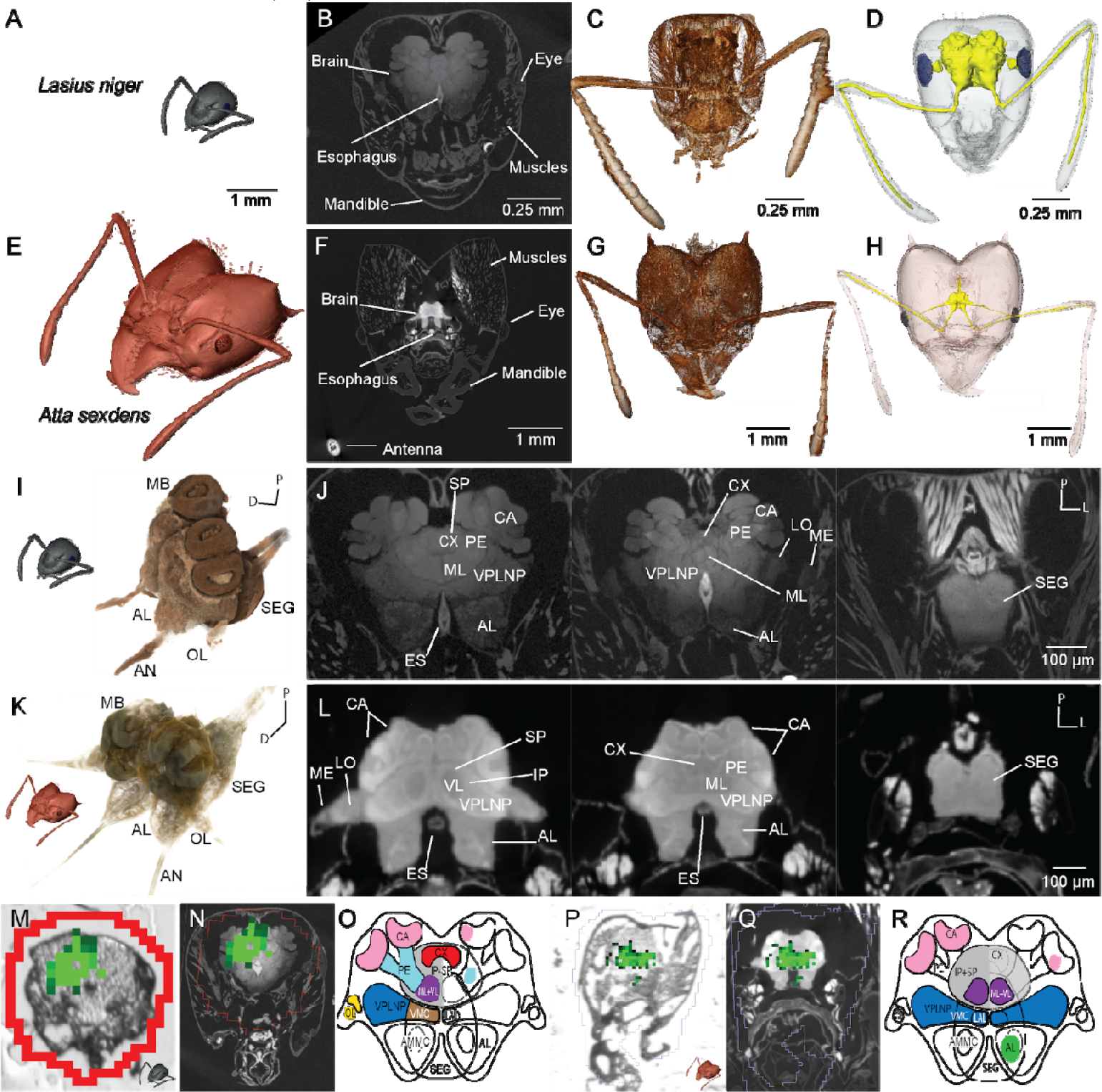
3-dimensional head anatomy of *L. niger* and *A. sexdens* with neuropeptide TK1 localization. (**A-H**) Anatomic proportions of brain to head ratios between *Lasius niger* (**A-D**) and Atta *sexdens* (**E-F**). (**A, E**) Comparable scale of two μCT based 3D models of two worker castes; (**B, F**) virtual plane of transverse sections through the 3D μCT volume; (**C, G**) Volume renderings (Drishti software version 2.6.4); (**D, H**) surface reconstructions of head capsule, brain, antennal nerves, and eyes (AMIRA™ software version 6.3). **I-L** Detailed brain anatomy resolved for *L. niger* (**I, G**) and A. *sexdens* (**K, L**) using μCT. **I, K** volume renderings of both brains; **G, L** virtual planes of transverse sections through the 3D μCT volume. **M-R** Co-registration of MALDI MSI ion map of neuropeptide TK1 for *L. niger* (**M-O**) and A. *sexdens* (**P-R**). **M, P** Ion map on a tissue section of TK1; **N, Q** overlay of MALDI MRMS MS and 2D μCT virtual slice; **O, P** brain drawing of the TK1 distribution of the tissue sections shown in M and R, respectively. AL, antennal lobe; AMMC, antennal mechanosensory and motor center; AN, antennal nerve; CX, central complex; ES, esophagus; IP, inferior protocerebrum; LAL, lateral accessory lobe; MB, mushroom bodies (CA, calyx; ML, medial lobe; PE, pedunculus; VL, ventral lobe); OL, optical lobe (LO, lobula; ME, medulla); SEG, subesophageal ganglion; SP, superior protocerebrum; VMC, ventromedial cerebrum; VPLNP, ventro-posterolateral neuropils. Orientation: D, dorsal; L, lateral; P, posterior.

Particularly challenging was to sufficiently contrast the brain and its substructures inside the intact head capsule, as the impermeable head capsule caused a blockage of the contrasting agents. Structures such as protective exoskeletons or shells present a persisting problem for staining solutions and are conventionally removed when working with arthropods or other invertebrate organisms (23). Notably, microdissection of such small specimens is time consuming, requires a high level of skill and can induce distortions up to major damage of the delicate micrometer-scale anatomy. Therefore, we severed the head from the body, allowing the phosphotungstic acid solution to diffuse directly through the nerve cord into the brain (**Suppl. Fig. S3**). Using this procedure, we were able to obtain very small volume pixel (voxel) sizes (*L. niger*: 0.805 μm, *A. sexdens*: 3.111 μm) at sufficient contrast and s/n to resolve fine structures of both brains. Moreover, not only brain and subesophageal ganglion were contrasted but off-branching nerves into antennae, eyes, mandibulae and maxillae of both species also showed high levels of contrast. Structures of different materials, less rich in lipids, such as musculature or cuticle were not penetrated equally strong by the contrasting solution. The strong contrasting of the nervous tissues enabled us to semi-automatically segment the brain and its surrounding tissues for subsequent 3D surface reconstructions (see **Fig. 3**).

The ant brain follows the general design of insect brains and, as in other holometabolous insects, is fused to the subesophageal ganglion (51). With the μCT dataset of each species, both high-resolution 3D anatomical models enabled us to navigate through the ant brains and the subesophageal ganglion to identify the organ’s subdivision. Following the nomenclature for brain proposed by Ito *et al*. (52) and Bressan *et al*. (53), we recognized 20 brain compartments in the μCT dataset of *L. niger* and 16 in *A. sexdens*, including optic lobes (OL), mushroom bodies (MB), central complex (CX), antennal lobes (AL), superior and inferior protocerebrum (SP+IP), antennal mechanosensory and motor center (AMMC), lateral accessory lobe (LAL), ventro-posterolateral neuropils (VPLNP), ventromedial cerebrum (VMC) and also the subesophageal ganglion (SEG) in both species (see **Fig 3I-L**). Due to the almost four-fold increased magnification for *L. niger* compared to the *A. sexdens* head, we could resolve more neuropils in *L. niger*, for instance lamina in OL and central body upper and lower units, protocerebral bridge and noduli in the central complex (**Suppl. Table S2**).

To date, ant brain neuropil maps were established by using confocal laser scanning microscopy (53, 54). Although this technique provides subcellular resolution and cell-specific staining, μCT does not require dissection of the brain, immunostaining followed by extensive imaging times and is therefore considerably less time-consuming. For instance, due to the specificity of the antibodies for synapsis detection, the brain map built by Bressan *et al*. (53) resulted in a highly detailed structure of the axon fascicles associated with compartment boundaries. Most of these structures were not observable in the here generated μCT atlas. However, methodologies have been developed to combine antibody labeling with heavy metal contrasting for generating a tissue-specific contrast detectable with μCT (37). Today, the increased imaging and processing speeds of high-resolution μCT provide a tool to quantitatively compare the neuroanatomy of many individuals across different developmental stages and castes to screen for heterogeneities in the plasticity of the brain.

For confocal- and/or laser scanning microscopy, the in-depth resolution (z-axis) is optically limited to about half of its lateral resolution (y, x-plane). This results in unequally shaped (anisotropic) voxels, 3D ‘volume pixels’, whereas μCT provides the same voxel edge length along all axes (isotropic voxels) (55). This allows for the calculation of distortion-free biovolumes that without interpolation allow for quantitative analyses. Both ant brain atlases generated here by μCT display the anatomical relationships between the brain, musculature and head capsule including mandibles and antennae (see **Fig.3**). Additional optical images of cryo-sections from snap-frozen samples served as controls for shrinking artefacts from the alcohol-based μCT contrasting protocol.

Despite nearly two orders of magnitude difference in head volume (*L. niger*: 0.031 mm^3^ and *A. sexdens*: 1.38 mm^3^), the brain of *L. niger* is only half the volume of *A. sexdens* (*L. niger*: 0.026 mm^3^ and *A. sexdens*: 0.052 mm^3^) (see **Fig. 3**). However, the ratio of brain volume as compared to the overall head volume was much higher in *L. niger* than in *A. sexdens*, revealing that the brain of *L. niger* consumes ∼80% of the head capsule whereas in *A. sexdens* only ∼7% of the head volume was occupied by the brain. These precise volumetric measurements provided valuable information when investigating the relationships between size and function of the brains. For example, once the distinct localization of a neuropeptide is known, quantitative LC-MS/MS in combination with μCT could in future studies allow for calculating the neuropeptide concentrations in each (sub)region of the brain and a comparison between different castes. Complementing the 3D microstructure in correlation to the 3D neuropeptide chemistry of the brain could shed light on how morphological adaptations, such as miniaturization, relate to behavior of ants and other social insects (56).

### Integrative spatial peptidomics and anatomy shows brain region specific chemistry

Integrating the serial spatial peptidomics maps into the anatomic μCT generated models resulted in 3D neuropeptide atlases of *L. niger* and *A. sexdens*. We visualized the co-registration between the 3D distributions of TK1, ITG-like neuropeptide and the μCT-based surface reconstructions of *L. niger* and *A. sexdens* (see **Fig. 4**).

**Figure 4.**
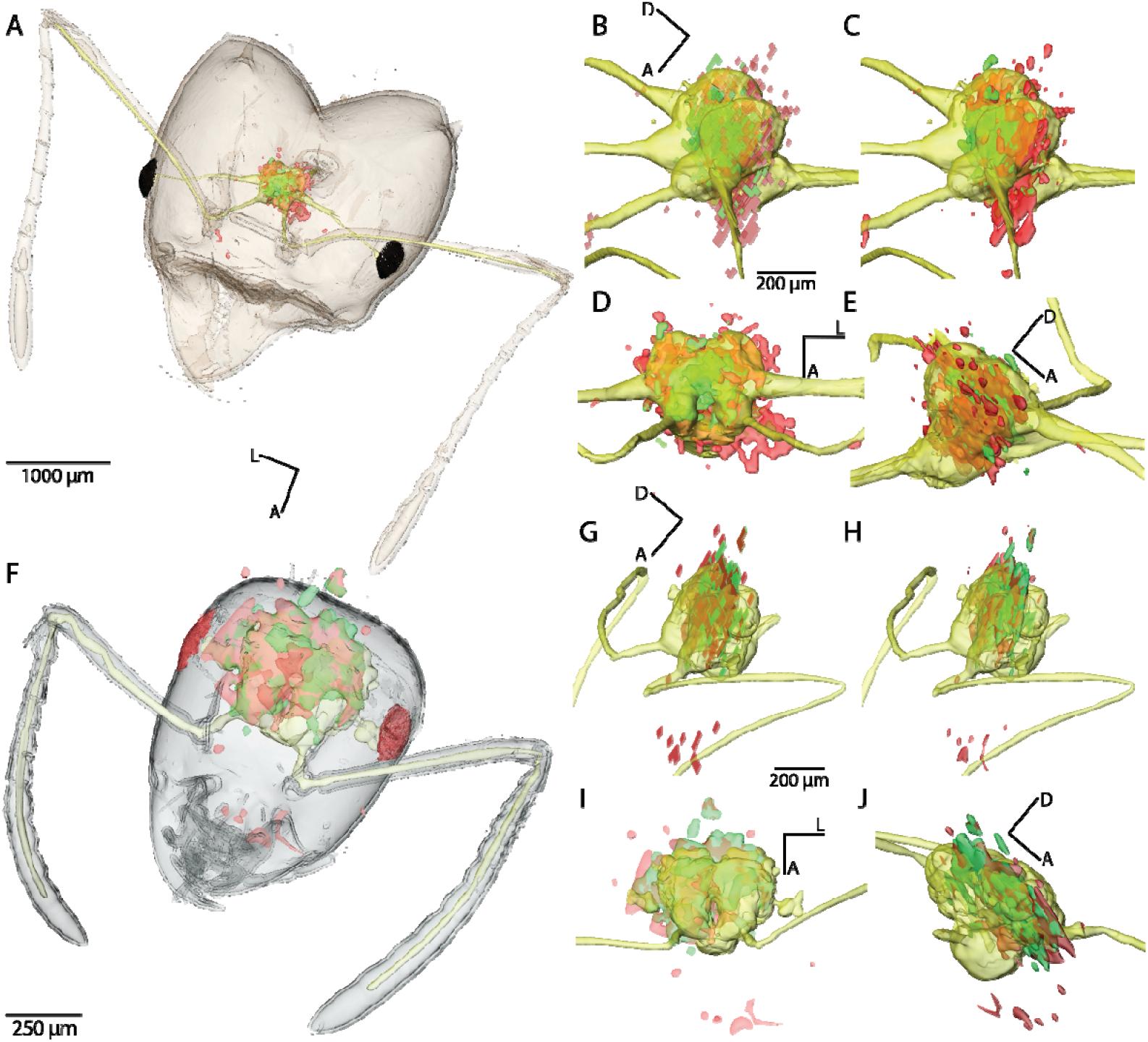
Integration of 3D neuropeptide distributions with micro-anatomy reconstructed 3D data. Surface renderings of segmented head capsule, brain/nerves and co-registered 3D distributions of TK1 (green) and ITG-like neuropeptide (red) in *A. sexdens* (**A**) and *L. niger* (**F**). **C-J**, Panels with four representations for surface reconstructions of the brain (yellow) and both neuropeptide distributions of each species (*A. sexdens*: **B-E** and *L. niger*: **G-J**); dorso lateral view, no smoothing and interpolation (top-left), dorso lateral view, smoothing and interpolation (top-right), dorsal view (bottom-left), latero-ventral view (bottom-right).

To create a reliable 3D co-registration, we based the matching between MSI and μCT data on manual annotations by experts in neuro- and invertebrate anatomy. The bright-field microscopy images of the tissue sections, used for MSI, provided key landmarks to determine in which orientation the chemical images had to be co-registered into the digital μCT datasets. There are three challenges for co-registering MSI data with other imaging modalities (27). Firstly, chemical distributions of analytes recorded by MSI must not necessarily correlate with a distinct anatomy of a tissue or organ. Secondly, MSI sensitivity for ion detection, the number of molecules that need to be ionized per pixel, limits MSI spatial resolutions to one or two orders of magnitude lower than radiation-based techniques, such as light microscopy (57). Thirdly, MSI is a surface-scanning technique that requires physical sectioning of frozen or chemically-fixed tissues, leading to distortions and subcellular damage of the tissue. Therefore, we based our co-registration between MSI and μCT on the optical images of the thaw-mounted cryo-sections that were recorded with a low-resolution slide scanner. Due to the generally limited quality of slide-scanner images, sub-regions of the brain were difficult to identify. Notably, we chose to use a slide scanner over an automated microscope, because it provided the fastest way to record the whole section series as swiftly as possible between cryo-sectioning and MSI to prevent oxidation of the lipid-rich brain tissue. For the co-registration, distinct tissue features and their relation to each other such as eyes, head capsule, esophagus, subesophageal ganglion and mushroom bodies served as reference points for a precise alignment of optical images, used for MSI and the corresponding μCT image plane (see **Fig. 3**; **Suppl. Fig. S4**). Critical for our workflow was to ensure that the correct proportions of the co-registered volumes, the z distances between the individual tissue sections were documented during sectioning and scaled proportionally during co-registration with μCT data (**Suppl. Fig. S4**). In a last step, once the optical images were individually matched to their corresponding virtual μCT image plane (see **Suppl. Fig. S4**) their coordinates in the 3D space could be transformed to the MSI neuropeptide maps and visualized as 3D volumes (see **Fig. 4**).

Although different animals for both MSI and μCT were used, we chose comparable castes and sizes to better correlate neuropeptide distributions with ant brain anatomy. The co-registration of chemical and anatomic data enabled us to calculate volumetric relations of brain size, relative amounts of neuropeptide and overall size of the head. For instance, TK1 and ITG-like neuropeptide occur in similar regions throughout the brains of *L. niger* and *A. sexdens*. Despite the substantially different anatomy and lifestyle of both ant species, their brain size is within the same order of magnitude and the distribution of essential neuropeptides, such as TK1 and ITG-like peptide, correlates between both species. Although conclusions from this comparison will have to be tested quantitatively, our comparative snapshot could help us to determine conserved and distinct features of the neurometabolism between ant species.

Even though MSI imaging provides an unprecedented view of neuropeptide distributions at anatomic resolution, exploration of the magnitude of chemistry detected has to be done. We provide the datasets, ready to browse online, for lipids and many other non-peptide metabolites in www.metaspace2020.eu (ant brain) as a community resource for further insect brain chemistry questions.

### Spatial peptidomics reveals neuropeptide distribution differs in ant brains

The region-specific localization of ten neuropeptides in both species strongly varied (**Table 2**). Consequently, we cannot exclude the presence of low neuropeptide levels in other regions of the brain. However, both species were measured with comparable settings allowing us the compare the spatial distribution of relative abundances of neuropeptides otherwise impossible with label-free omics approaches. In general, neuropeptides of *L. niger* showed wider distributions compared to *A. sexdens*. This could reflect *L. niger* being a species with only one morphological caste of workers with less specialization than in *A. sexdens* where at least six different kinds of specialized workers operate delimited tasks (58). Regarding the occurrence of the neuropeptides in different brain regions, TK1, ITG-like and sNPF appeared widely distributed in the neuronal organ, exhibiting a similar brain neuropil allocation in both species (see **Table 2/Fig. 4**). Similarly, NPLP1-4 and NPLP1-6 were detected across most regions of the brain of *L. niger*, but only in the ventro-posterolateral neutropil region of *A. sexdens*. TK4 and IDLSRF-like peptide were imaged in half of the brain regions annotated for *A. sexdens* and found in most regions across the brain of *L. niger*. The co-registration between both techniques that were applied to different individuals restricted an allocation of neuropeptides to the level of the annotated brain regions. However, the confidence of identification and co-registration per brain region was increased due to our 3D approach, as we could trace each region throughout the consecutive sections of our serial brain slices.

**Table 2.**
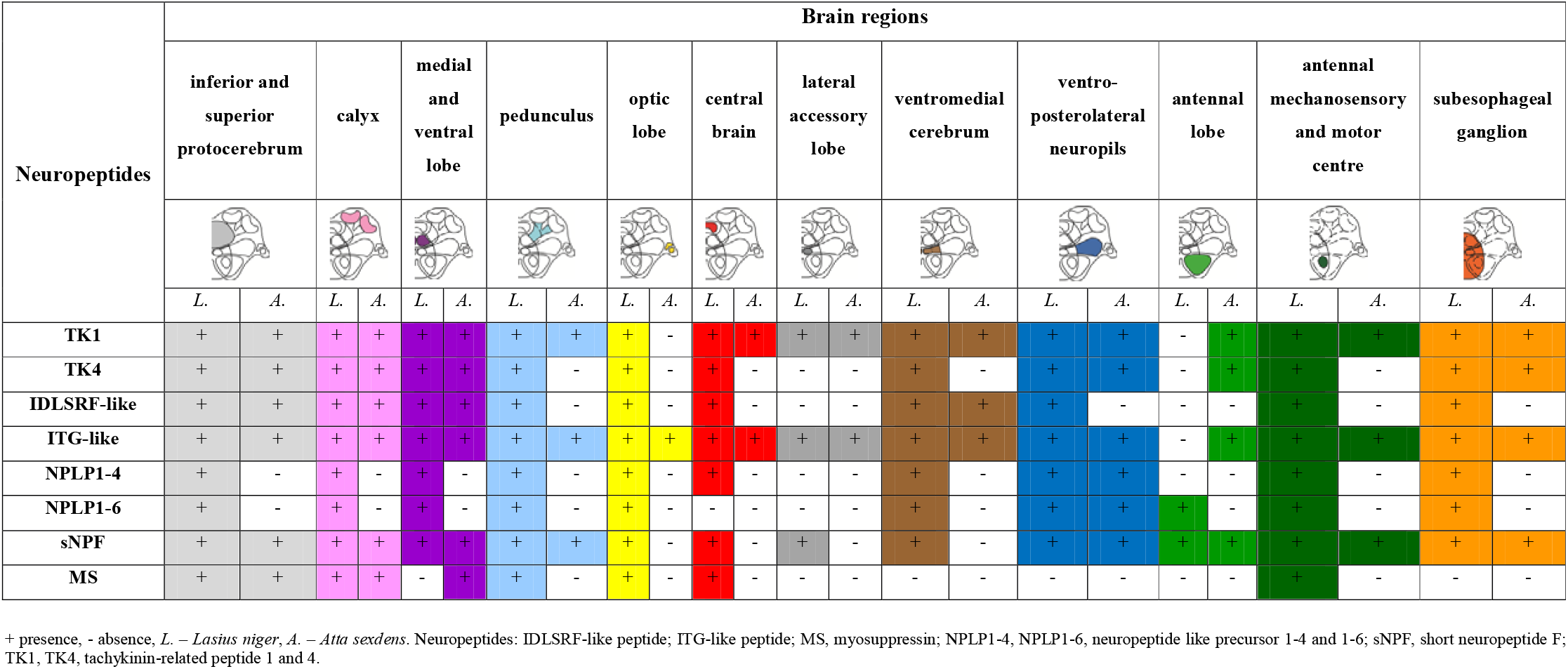
Distribution of the common neuropeptides detected and identified from brain tissue sections via spatial peptidomics after co-registration with μCT data set in *L. niger* and *A. sexdens*.

How neuropeptide distributions, for example left-right asymmetries, regulate each other and relate to neurobiological processes such as lateralization and ultimately behavior remains to be determined. Considering that neuropeptide distributions vary along the animals’ lifecycles, these changes can be very fast for some developmental stages. For instance, in the honeybee (*Apis melifera*), two neuropeptides change significantly during the first 25 days of the adult stage (22). Validating our findings through subsequent studies of larger cohorts with animals of different castes and developmental stages could reveal how spatial heterogeneity of neuropeptides in correlation to microanatomic brain plasticity shape social behavior in ants and other insects.

### Atlases with multi-omics data are emerging tools for organismal biology

Applications of multi-omics 3D atlases that combine anatomic structure with spatial proteomics, transcriptomics or metabolomics data has become an emerging field (26, 27, 59, 60). These modern multi-omics atlases build upon the extensive knowledge generated from large-scale descriptive studies that for instance mapped the 3D ultrastructure of a whole fly brain (61), or the early human development imaged by light-sheet microscopy (62), by adding a layer of functional data – a combination of structure and function. Still, despite rapid improvements on the technological side in terms of imaging speeds, spatial resolution, data processing and cost (16), there are still major challenges making large-scale correlative studies difficult. Besides harmonization of sample processing workflows for both 3D imaging techniques, the correlative analysis between modalities remains the most critical step. Co-registration (63, 64) of the different types of imaging data and the correlative analysis across modalities (65) has not yet been integrated into a user-friendly software with a graphical user interface. Spatial omics techniques, such as MSI, are often multiplexed and enable the detection of hundreds to thousands of signals at once without prior knowledge of the sample. Although the data are more complex and the analysis challenging, untargeted analysis comes with the unique advantage that it allows for the discovery of hitherto unknown biological mechanisms if known anatomic or cellular structures are correlated to spatial chemistry (66, 67).

In this study, we highlight the power of spatial peptidomics for deciphering ant neurochemistry. The provided insights into key signaling molecules ultimately allows for identification of factors that drive caste identity and behavior. Our approach offers an integrative 3D visualization of the native, label-free biochemistry with undisturbed microanatomy allowing for comparisons of neuropeptide levels across the different subregions of the brains of two ant species. Our atlas contributes to a better understanding of the processes and interactions between signaling molecules in the brains of ants and other insects that have allowed for their high level of coordination to not only thrive in nearly every ecosystem but to become essential for ecosystem health.

### Material and Methods

#### Specimen

Workers of the monomorphic black garden ant *Lasius niger* were taken from laboratory colony maintained in an incubator in summer conditions (14 h light, 27°C/ 10 h darkness, 20°C; 60% humidity) and fed *ad libitum*. Medium and large castes of polymorphic leafcutter ant *Atta sexdens* were collected from Schönbrunn Zoo colony (Vienna, Austria).

#### Genome search and mass calculation of neuropeptides

Published neuropeptide precursor sequences of *Acromyrmex echinatior* (45) and *Camponotus floridanus* (46) were used to query the NCBI Whole-Genome Shotgun (WGS) database of *Atta cephalotes* (project number PRJNA279976) (48) and *L. niger* (project number PRJNA269328) (47) via tBLASTn online using BLOSUM62 matrix. Hits with the lowest E-values were manually annotated and the peptide sequence was determined according to the obtained alignment of the query and the subject. Peptide masses were calculated using Peptide Mass Calculator v3.2 online tool (http://rna.rega.kuleuven.ac.be/masspec/pepcalc.htm).

#### RNA isolation and sequencing of myosuppressin, tachykinin-related peptide, allatostatin and short neuropeptide F precursors

RNA extraction of 2-5 heads of *L. niger* or *A. sexdens* was done using Quick-RNA™ MiniPrep kit (Zymo Research). Samples were homogenized in 350 μl Lysis Buffer with 2-4 BashingBead™ (Zymo Reseach) using homogenizer Precellys 24 (Peqlab) for 3 × 30 sec 6000 rpm. DNase I treatment was performed after eluting RNA from columns prior to reverse transcription accordingly: samples (50 μl) in 0.5x DNase I buffer (Thermo Scientific, #B43) were incubated with 1 U of DNase I for 20 min at r. t. following the inactivation of DNase I by adding EDTA to 2.5 mM final concentration and heat denaturation at 70°C for 10 min. Reverse transcription was performed using High-Capacity cDNA Reverse Transcription Kit (Applied Biosystems) according to manufacturer’s instructions. PCR products of precursor sequences were obtained via standard PCR using Phusion Hot Start II DNA Polymerase (Thermo Scientific) and primers (Sigma-Aldrich) (**Suppl. Table S3**). The PCR fragments were cut from an agarose gel, purified using GeneJET Gel Extraction kit (Thermofisher Scientific) and sequenced at LGC Genomics (Berlin, Germany) using both reverse and forward primers. Using standard PCR we obtained only the beginning (∼700 bp) of *A. sexdens* tachykinin-related peptide precursor, while the end was confirmed via Rapid Amplification of 3’ cDNA Ends (3’ RACE) technology: reverse transcription was performed using 3’-Race-RT-primer, first PCR was done using 3’-Race-PCR-Rev and A.sex-TK-Fw primers and then the product of nested PCR (361 bp), obtained using 3’-Race-PCR-Rev and 3’-Race-PCR-Fw primers and diluted reaction of first PCR as a template, was sequenced. The confirmed *A. sexdens* TK precursor sequence is much shorter compared to *A. cephalotes* or *A. echinatior* (**Suppl. Fig. S1**). The sequence of *L. niger* TK precursor was confirmed by obtaining and sequencing two overlapping PCR products using primers L.nig-TK-Fw1+L.nig-TK-Rev1 for the N-termini and L.nig-TK-Fw2+ L.nig-TK-Rev2 for C-termini.

#### MALDI MRMS imaging and data analysis

Animals were placed in a reaction tube and killed by immersion in liquid nitrogen for 5 sec (*L. niger*) or by freezing at −80°C (*A. sexdens*) and decapitated under the stereomicroscope. Heads were immersed in a mold with 10% gelatine (cold water fish skin, SIGMA 67041) to obtain a block suitable for cryo-sectioning at 20°C (Leica CM1950). Dorso-ventral serial sections of 10 μm were mounted onto conductive indium-tin oxide (ITO)-coated glass slides (Bruker Daltonics GmbH & Co KG, Bremen, Germany).

Slides were dried under vacuum for 15 min and then coated with alpha-cyano-4-hydroxycinnamic acid (HCCA) using an ImagePrep device (Bruker Daltonics, Bremen, Germany). HCCA matrix (7 mg/ml dissolved in 50% [v/v] acetonitrile, 0.2% [v/v] trifluoroacetic acid) was sprayed with the ImagePrep default program for this matrix in five phases. In the first initialization phase, matrix was applied in 10 cycles with a fixed spray time of 2.5 sec, spray power set to 15, modulation 40, incubation time 15 sec, and dry time 65 sec. In phases 2–5, spray time as well as dry time were automatically controlled by the instruments integrated scattered light sensor. The spray power was set to 25, modulation 45, and incubation time was 30 sec.

Data was acquired on a Bruker solariX XR MRMS instrument (Bruker Daltonics, Bremen, Germany) equipped with a 7 T superconducting magnet, MALDI ion source and smartbeam II laser technology. Compass ftmsControl was used to set the instrument parameters. 1000 laser shots were collected per spectrum in the mass range of 300-3000 Da (*L. niger*) or 300-2600 Da (*A. sexdens*). The raster size was set to 30 μm (*A. sexdens*) or 40 μm (*L. niger*) resulting in datasets containing 143359 spectra or 25707 spectra, respectively. Mass calibration was performed externally on sodium trifluoroacetate in ESI mode. The mass resolution was greater than 220,000 at m/z 400. FlexImaging Software version 4.1 was used to define the measurement areas. Raw data for each ant was imported to the software SCiLS Lab version 2023a (Bruker Daltonics, Bremen, Germany) and normalized to total ion count (TIC). For the 3D MALDI model, low resolution scan images were stacked virtually in z-direction according to the section outline with 10 μm distance using the SCiLS Lab registrator. Neuropeptide distributions in 2D and 3D were visualized by false-color coding and low intensity pixel were made transparent.

#### μ-CT scanning

Specimens were placed 30 sec in a petridish on ice before decapitation with a sharp razor blade. The heads were fixed in Bouin’s solution overnight. Then the fixative was removed with disodium hydrogen phosphate buffer (0.1 M, pH 7.2, 1.8% sucrose) by 2 × 30 min washes, followed by an overnight immersion in the same buffer and 2 × 30 min washes next day. Afterwards, the heads were dehydrated in a graded ethanol series and contrasted for 2 weeks using a solution of 1% phosphotungstic acid in 70% ethanol, submersed in acetone and critical point dried (CPD) using a Leica EM CPD300 (Leica Mikrosystems GmbH, Wetzlar, Germany). Contrasting and CPD protocols were adapted from previous studies (35, 36, 38).

After CPD the heads were carefully mounted on the tip of a glass syringe using a hot glue gun. A nanotom m (GE Measurement & Control, Wunstorf, Germany), equipped with an X-ray cone beam setup was used for scanning. Magnifications were defined through geometrical relations that based on the distances between sample, source and detector. The heads were scanned at a voltage of 110 kV, 120 μA current and 0.75 s exposure time at an averaging of 4, which resulted in a scanning time of 90 minutes and 1500 number of projections acquired during scan. The tomographic reconstruction was performed using the standard parameters offered by the phoenix datos|x 2.2 reconstruction software on a separate workstation. *L. niger*: resulted in a voxel size of 0.805 μm and *A. sexdens*: 3.111 μm). Being able to move the smaller head of *L. niger* closer to the X-ray source and further away from the detector resulted in a higher magnification due to the X-ray cone beam. For feasible data processing, the 16-bit volume was cropped, converted to 8 bit depth and formatted to the *.hdr format in the 3D-visualization software VGStudio (2.2).

#### 3D visualization of μ-CT derived anatomy

3D volume exploration and rendering were done with the v3D volume exploration and visualization software Drishti, **v**ersion 2.5.1 (68) and Amira 6.3 (Thermo Fischer Scientific, USA). Segmentation and surface reconstruction of specific anatomical structures such as head capsule, eyes, brain and nerve branches were done in Amira v.6.3 in an adapted workflow after Ruthensteiner et al. (69).

#### Co-registration of μ-CT and MALDI-MSI data

MALDI-MSI and μCT datasets were combined in 3D space using the visualization software Amira v.6.3 (FEI Co., USA). The co-registration workflow had to be adapted as both MALDI-MSI and μCT datasets originated from different specimens and imaging techniques. Because the ant heads are heterogeneous tissue samples and were neither chemically fixed nor infiltrated with embedding medium, the cryo-sections showed strong distortions and ruptures. Therefore, orientation and subsequent fitting of the optical image stack into the μCT volume was based on visually comparable structures such as eyes, brain regions and mouthparts.

For the co-registration, the SCiLS aligned image stacks of the data were used. The general co-registration workflow consisted of two main steps. In the first step, the series of optical images of the MALDI-MSI sections was co-registered to the μCT data. In the second step, the same transformation parameters were then applied to the image stacks of the ion maps of TK1 and ITG-like neuropeptide. The same workflow was used for both ant species.

Initially, the Z-distances between the optical images had to be adjusted because not every section was used for MALDI-MSI. Gaps in the image stacks were filled with empty images. The optical image series of the tissue sections with the correct spacing was then imported into AMIRA. The corresponding sectioning planes between the optical images and μCT slices could be determined because the μCT volume was virtually sliced along each of the xyz-planes. With the series of optical images being aligned and the sectioning plane determined, the volume of the optical images was fitted into the 3D space of the μCT volume through manual translation, rotation and proportional scaling by aligning anatomical structures.

The ion image stacks of TK1 and ITG-like neuropeptide were then co-registered into the volume, using the same transformation parameters, determined with the optical image series. For visualization purposes, ion distribution stacks were also threshold-segmented and visualized as surfaces in the 3D model. The surface reconstructions allowed for simultaneous visualization of the ion maps in the 3D model based on different transparency settings.

## Data availability

All microscopy and micro-CT datasets can be directly downloaded from Figshare: (under process). MALDI-MSI data were deposited within a project on www.metaspace2020.eu database (link) and can be browsed online for lipids and other metabolite distributions.

## Acknowledgements

We thank Schönbrunn Zoo (Vienna, Austria) for *A. sexdens* specimen, Thomas Haider (Medical University of Vienna) for scanning the histological sections, Thomas Eder (University of Veterinary Medicine, Vienna) for help with sequence mining, Markus Gold-Binder (Medical University of Vienna) and Estela Pérez Santamarina (Medical University of Vienna) for technical assistance with cloning. We also thank Nikolaus Leisch (MPIMM, Bremen, Germany) for support with critical point drying and Bernhard Ruthensteiner (ZSM, Munich, Germany) for assistance during μCT procedures. We thank Jens Fuchser (Bruker) for help with MALDI imaging data recording. Work in the laboratory of C.W.G has been supported by the Vienna Science and Technology Fund (WWTF) through project LS13-017, the Austrian Science Fund (FWF) through project P32109, and the Ministry for Education, Science and Research of the Austrian Government through a ‘Hochschulraum-Strukturmittel-Projekte (HRSM)’. C.W.G and J.V.B received funding via a joint FWF and the Research Foundation Flanders (FWO) project (I3243). B.G and M.L would like to thank the Max Planck society for the financial support.

## Author contributions

E.G.M performed cryo- and histological sections and all manipulations with alive ants; Z.L searched the genomes and cloned the precursors, J.O and M.L performed MALDI imaging analyses; B.G carried out μCT sample preparation, measurements, image processing of μCT and MALDI-MSI data for 3D co-registration based on annotations by E.G.M and visualization. R.H, J.V.B and C.W.G analyzed data. J.O, M.L and C.W.G designed the study. All authors wrote the manuscript.

## Conflict of interest

J.O is employed at Bruker Daltonics GmbH & Co. KG, a vendor for mass spectrometry instruments and solutions.

## Supplementary Information

**Supplementary Figure S1.**
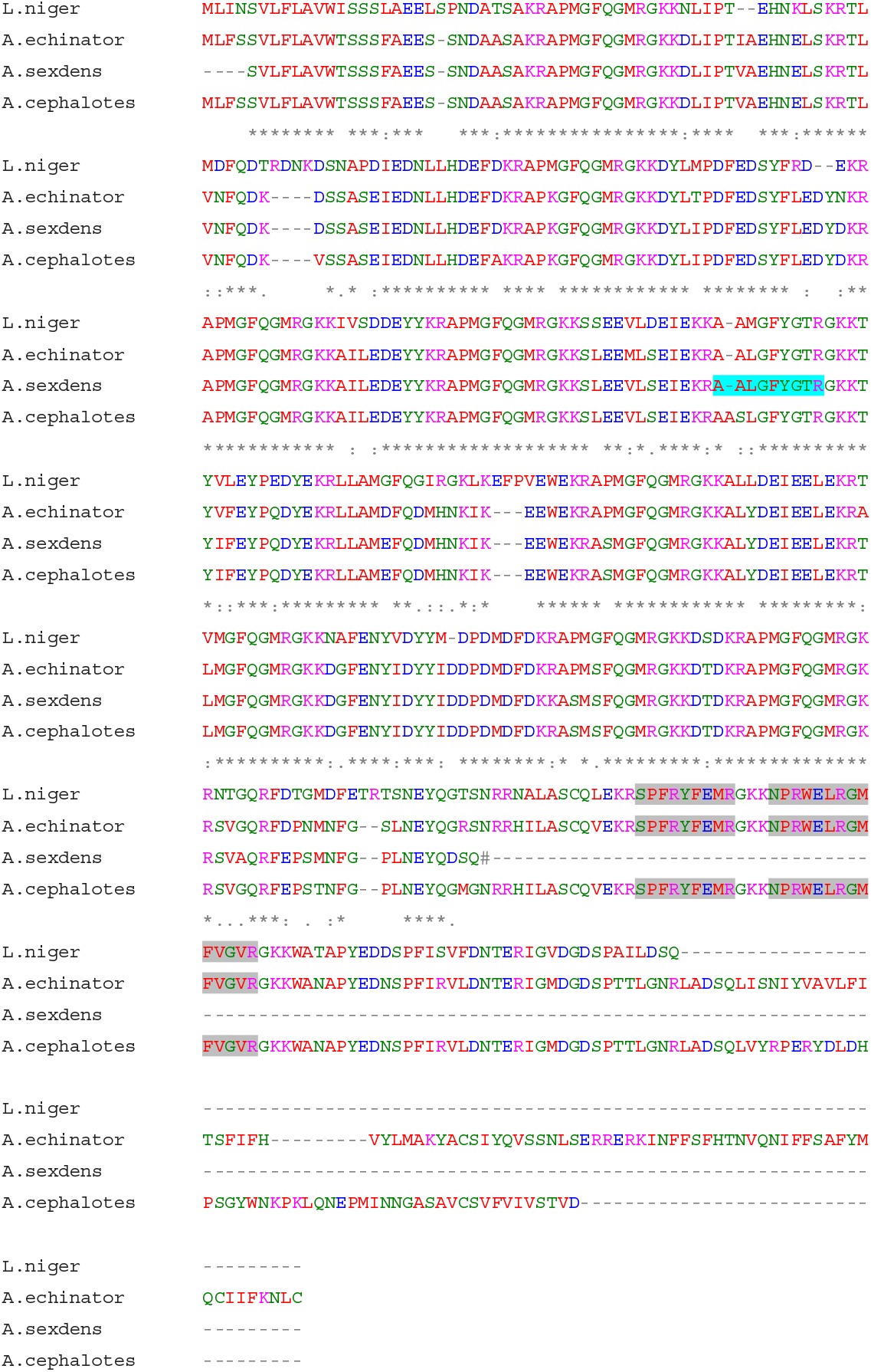
Alignment of tachykinin-related peptide (TK) precursors of *A. sexdens, A. cephalotes, A. echinator* and *L. niger*. *A. sexdens* TK7 was confirmed to be a single amino acid shorter compared to *A. cephalotes* (shown in light blue). The cloned *A. sexdens* TK precursor is shorter (stop codon indicated by #; using 3’ RACE and poly dT primer, see Suppl. Table S3) as compared to the other species lacking two neuropeptides (highlighted in grey, TK5 and TK6 according to the Suppl. Table S1); it needs to be confirmed whether a longer version co-exists.

**Supplementary Figure S2.**
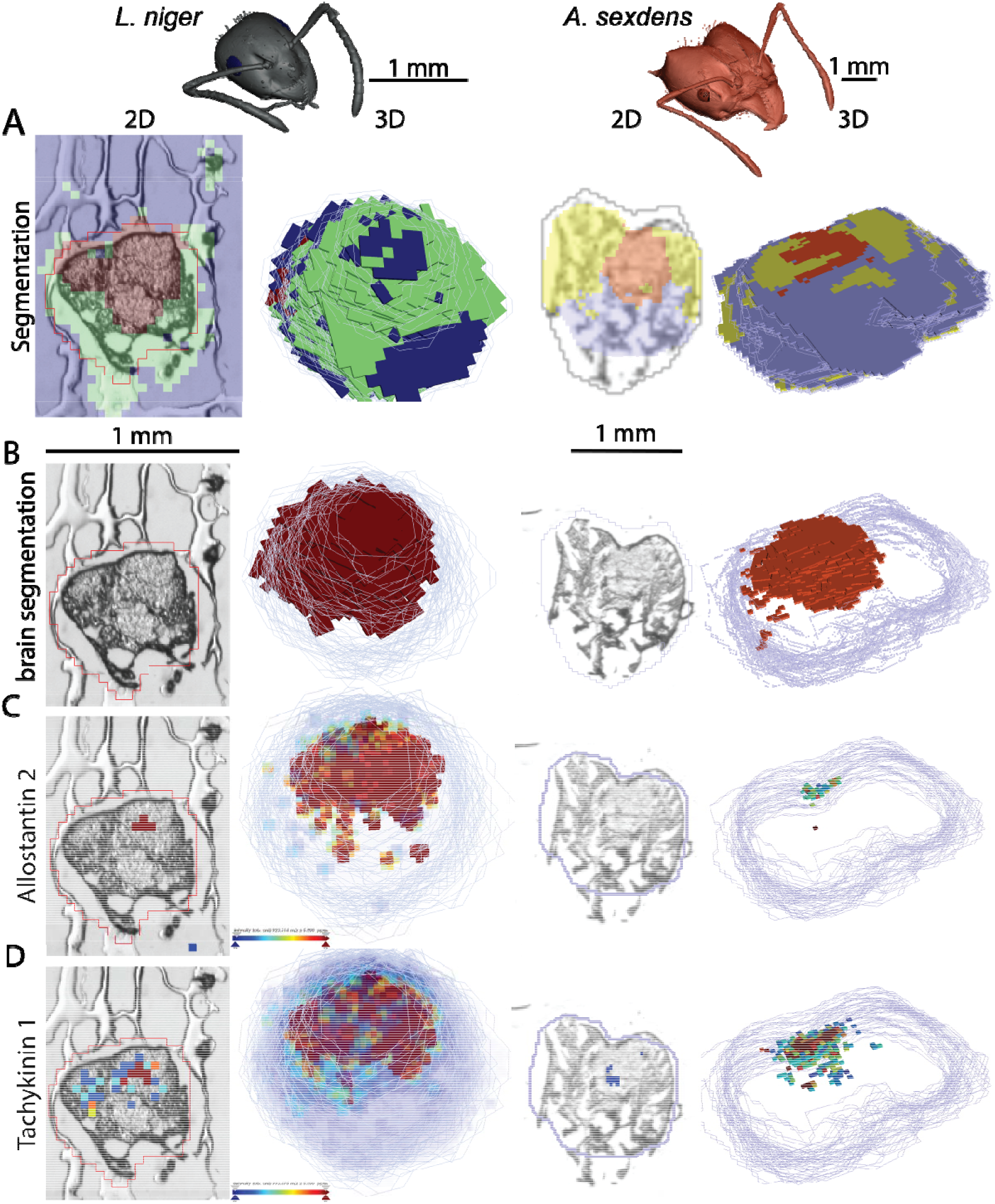
Spatial clustering of MALDI-MSI data acquired from section series of *L. niger* and *A. sexdens* heads. (**A**) Dataset of each head divided into three spatial clusters of which one was distinctly colocalized with the brain (red) centered in the head capsule shown in (**B**). Tissues surrounding the brain of *L. niger* clustered within the green cluster and in *A. sexdens* summarized within a yellow and purple cluster. In *L. niger* off-tissue signals were assigned to the blue cluster. (**C**) and (**D**) show heat maps of the peptide distributions for allatostatin (AstA) 2 and tachykinin-related peptide 1 (TK1) for a 2D plane and the 3D volume for each ant, illustrating the spatial heterogeneity along depth.

**Supplementary Figure S3.**
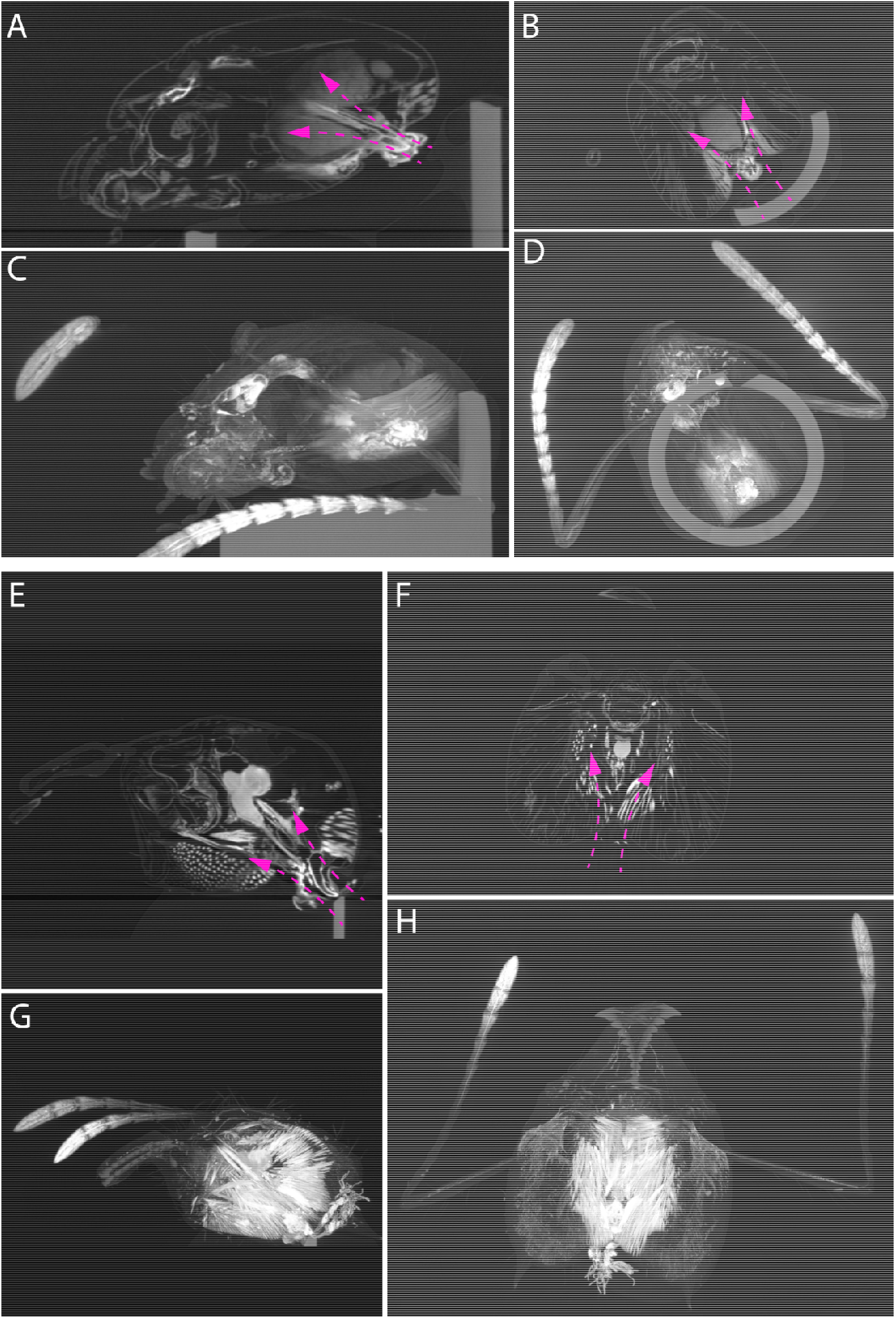
Penetration of the contrasting agent into the separated head capsules through the brain of *L. niger*. (**A-D**) **and *A. sexdens*** (**E-H**). Magenta arrows indicate direction of diffusion. (**A, B, E, F**) show 2D planes and (**C**,**D**,**G**,**H)** show maximum intensity projections along the z-axis of the 3D volumes.

**Supplementary Figure S4.**
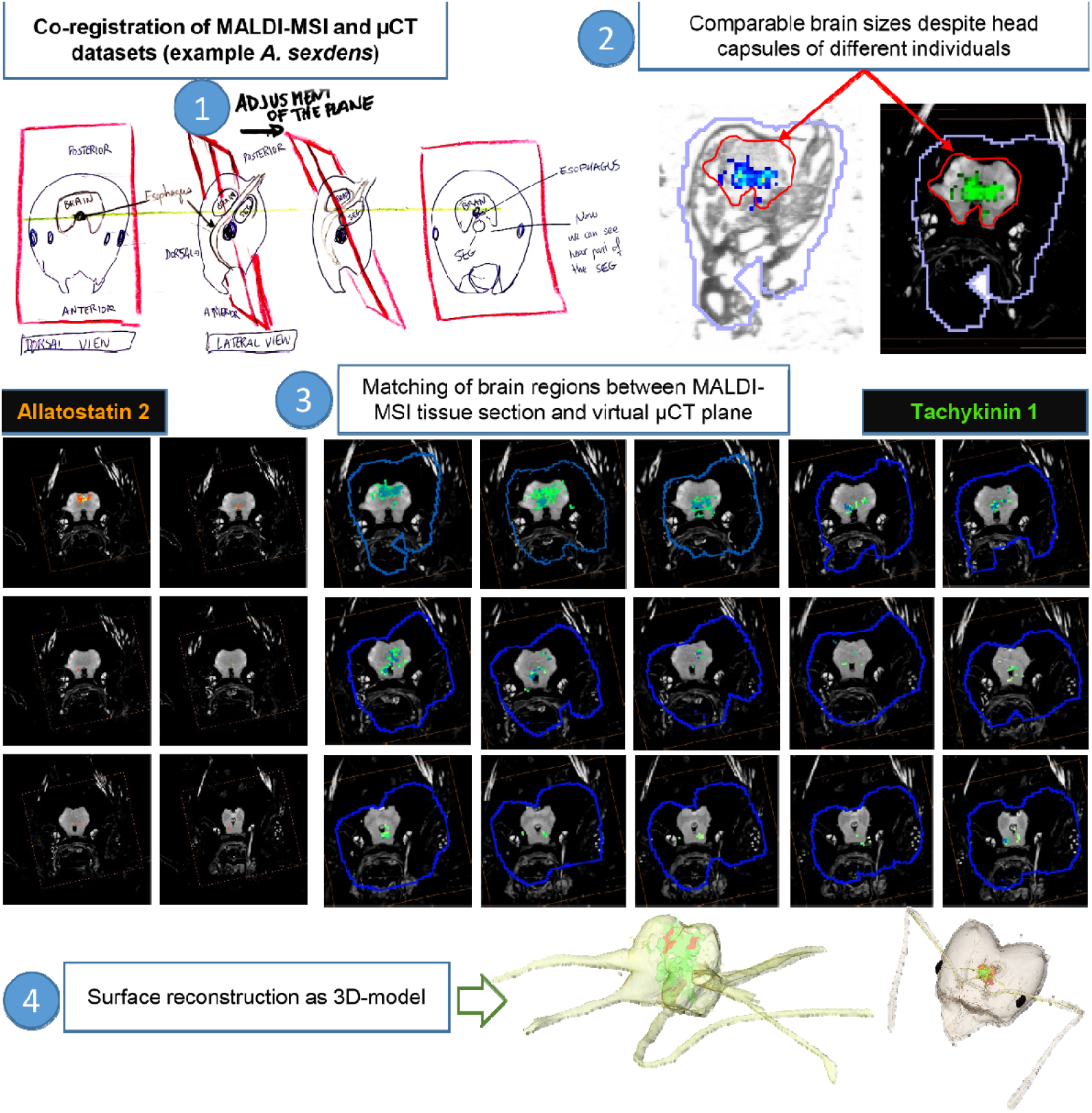
Co-registration between mass spectrometry imaging and microtomography data. Workflow consisted of four main steps: 1. Matching the virtual plane in the μCT data to the physical sectioning plane of the tissue sections, 2. Allocate brain regions, visible in both modalities (μCT and bright-field microscopy sections used for MALDI-MSI), 3. Align MALDI-MSI data with corresponding μCT slice based on bright-field microscopy image in 3D space (using Amira software), 4. Create surface renderings of each neuropeptide measured with MALDI-MSI to colocalize and simultaneously visualize within 3D microanatomy of each animal.

**Supplementary Table S1.**
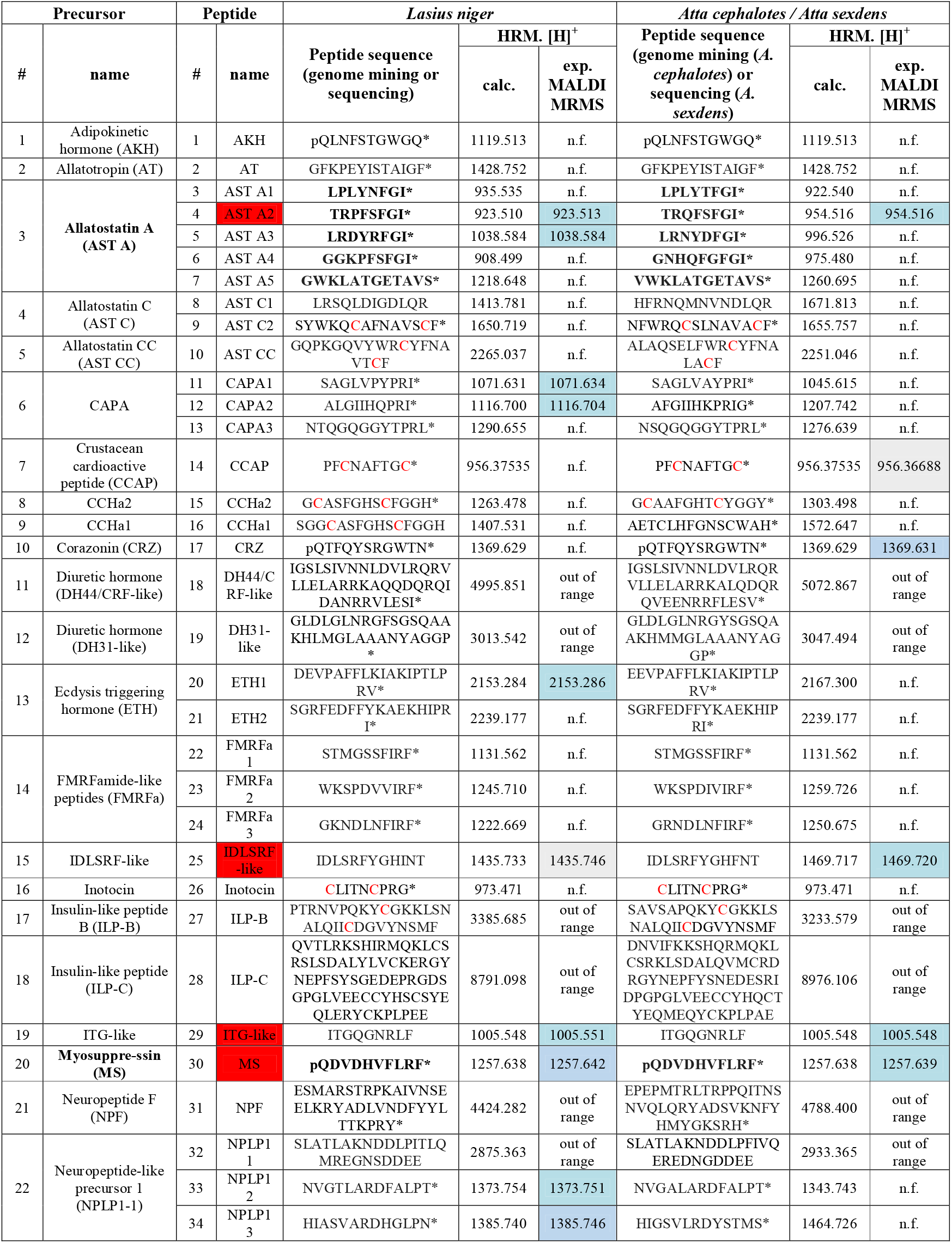
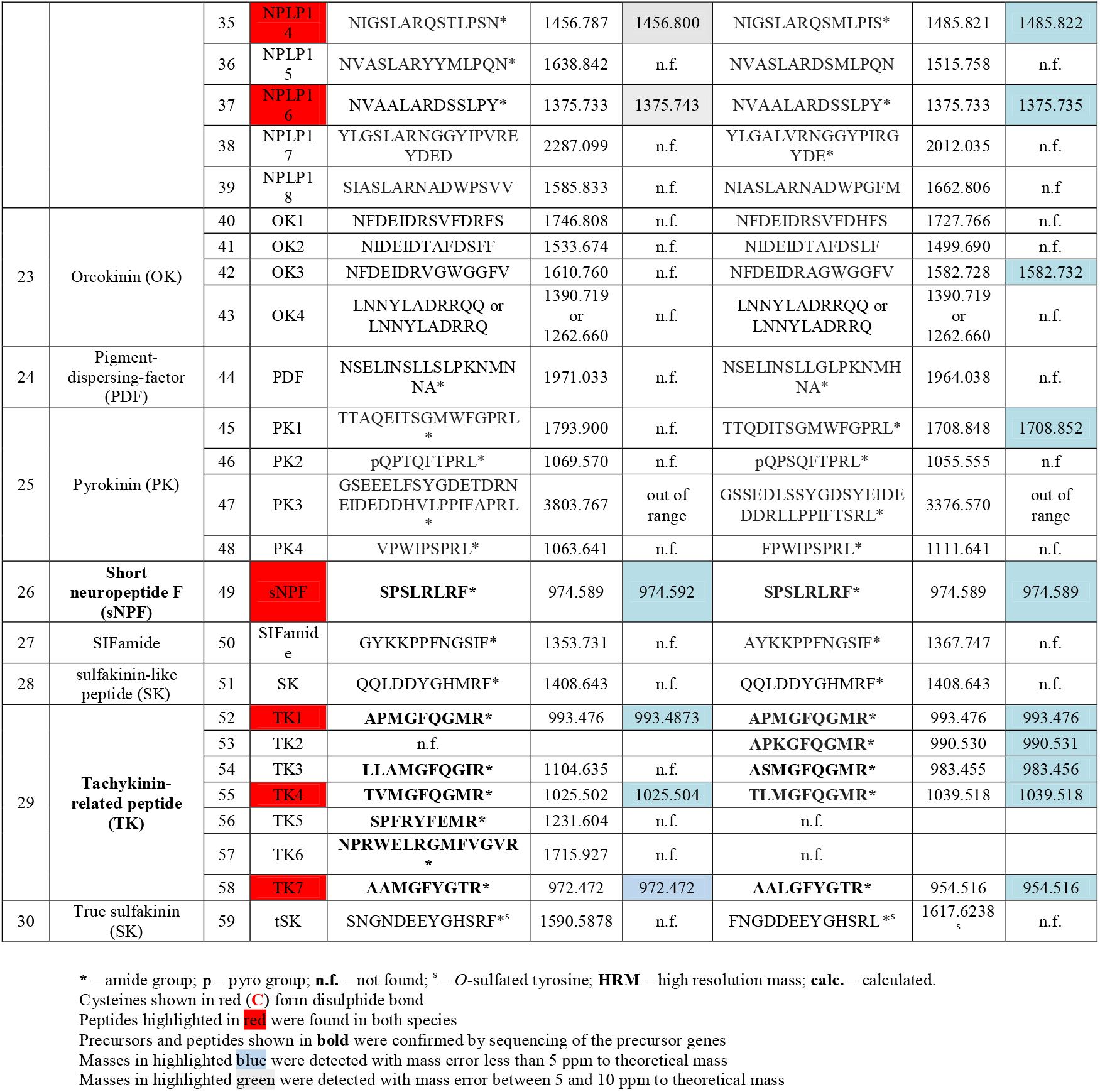
A summary of *L. niger* and *A. cephalotes/A. sexdens* neuropeptide sequences, masses and MALDI MRMS results.

**Supplementary Table S2.**
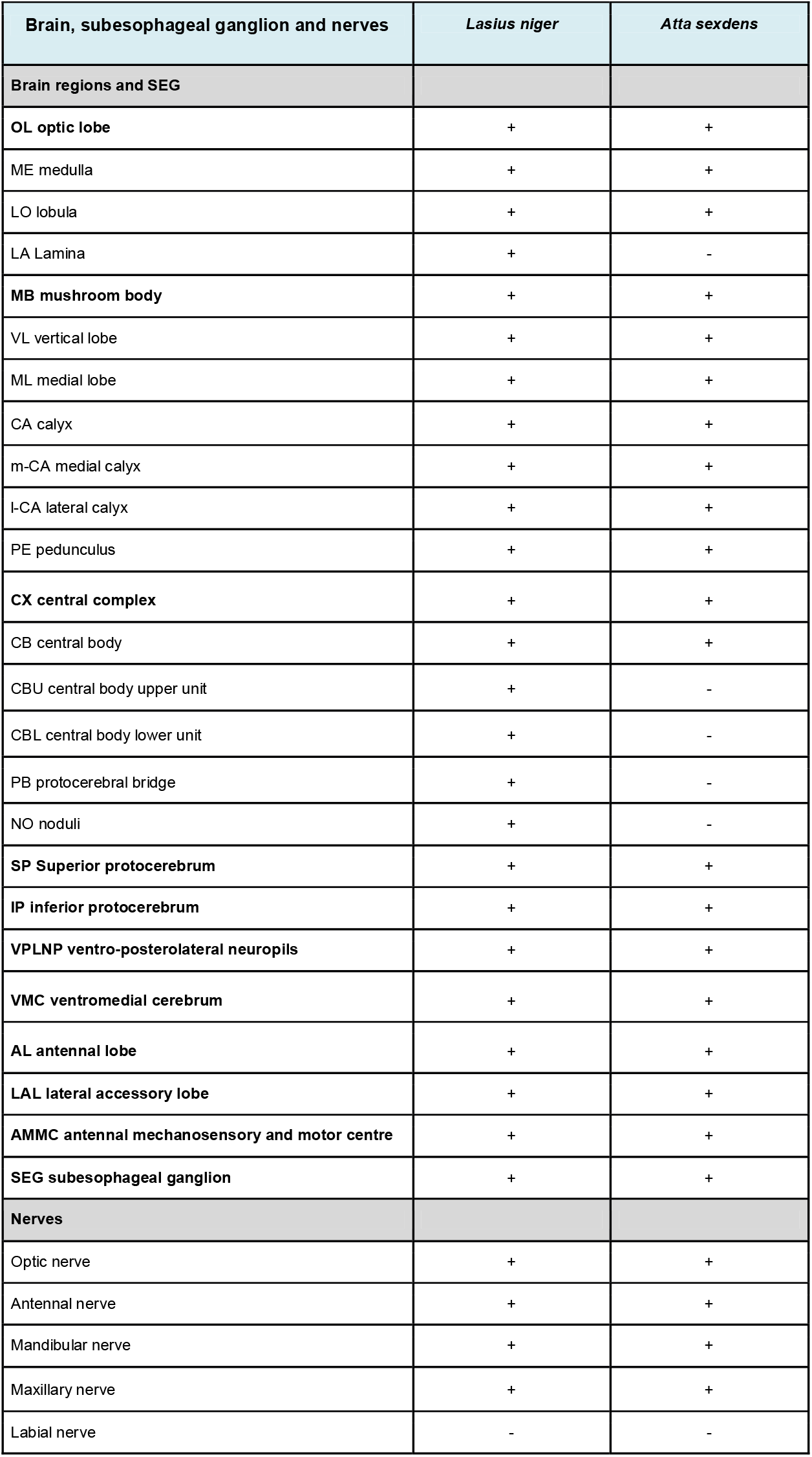
Resolved identified brain areas in *L. niger* and *A. sexdens* by using μCT.

| Brain, subesophageal ganglion and nerves | *Lasius niger* | *Atta sexdens* |
| --- | --- | --- |
| **Brain regions and SEG** |  |  |
| **OL optic lobe** | + | + |
| ME medulla | + | + |
| LO lobula | + | + |
| LA Lamina | + | - |
| **MB mushroom body** | + | + |
| VL vertical lobe | + | + |
| ML medial lobe | + | + |
| CA calyx | + | + |
| m-CA medial calyx | + | + |
| l-CA lateral calyx | + | + |
| PE pedunculus | + | + |
| **CX central complex** | + | + |
| CB central body | + | + |
| CBU central body upper unit | + | - |
| CBL central body lower unit | + | - |
| PB protocerebral bridge | + | - |
| NO noduli | + | - |
| **SP Superior protocerebrum** | + | + |
| **IP inferior protocerebrum** | + | + |
| **VPLNP ventro-posterolateral neuropils** | + | + |
| **VMC ventromedial cerebrum** | + | + |
| **AL antennal lobe** | + | + |
| **LAL lateral accessory lobe** | + | + |
| **AMMC antennal mechanosensory and motor centre** | + | + |
| **SEG subesophageal ganglion** | + | + |
| **Nerves** |  |  |
| Optic nerve | + | + |
| Antennal nerve | + | + |
| Mandibular nerve | + | + |
| Maxillary nerve | + | + |
| Labial nerve | - | - |

**Supplementary Table S3.**
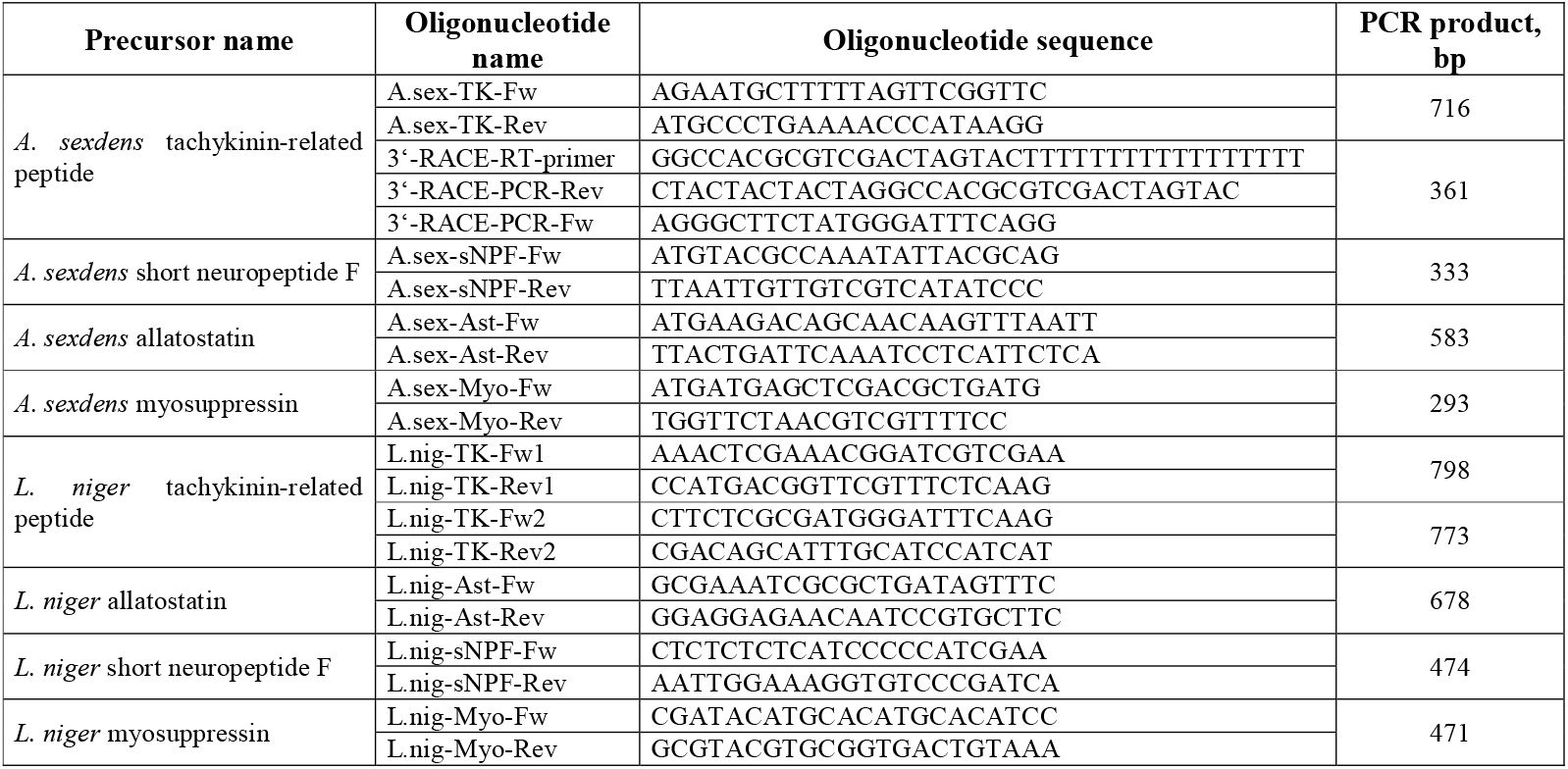
DNA oligonucleotides used in for PCR and sequencing of neuropeptide precursors.

**Supplementary Information S1.**
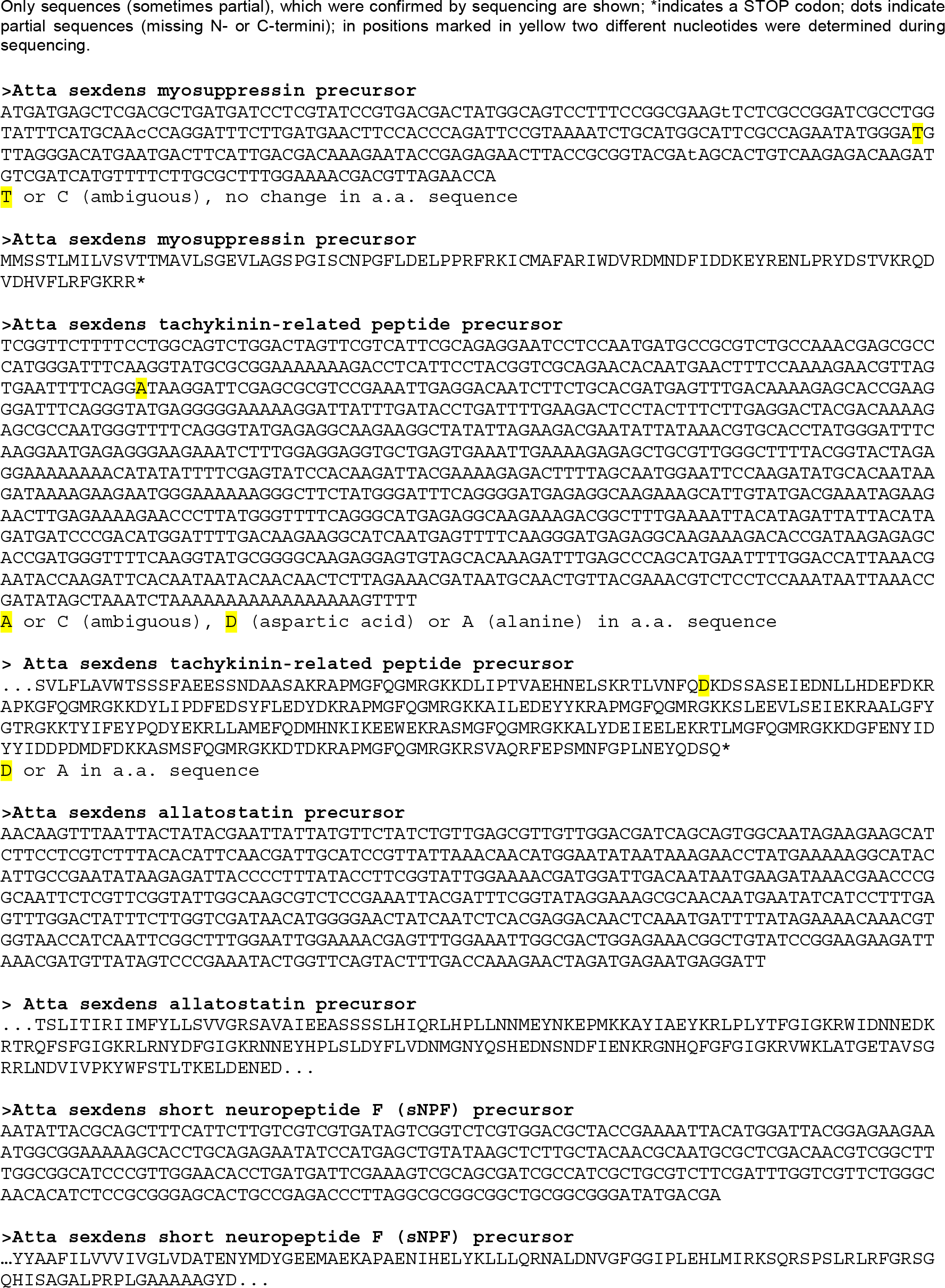
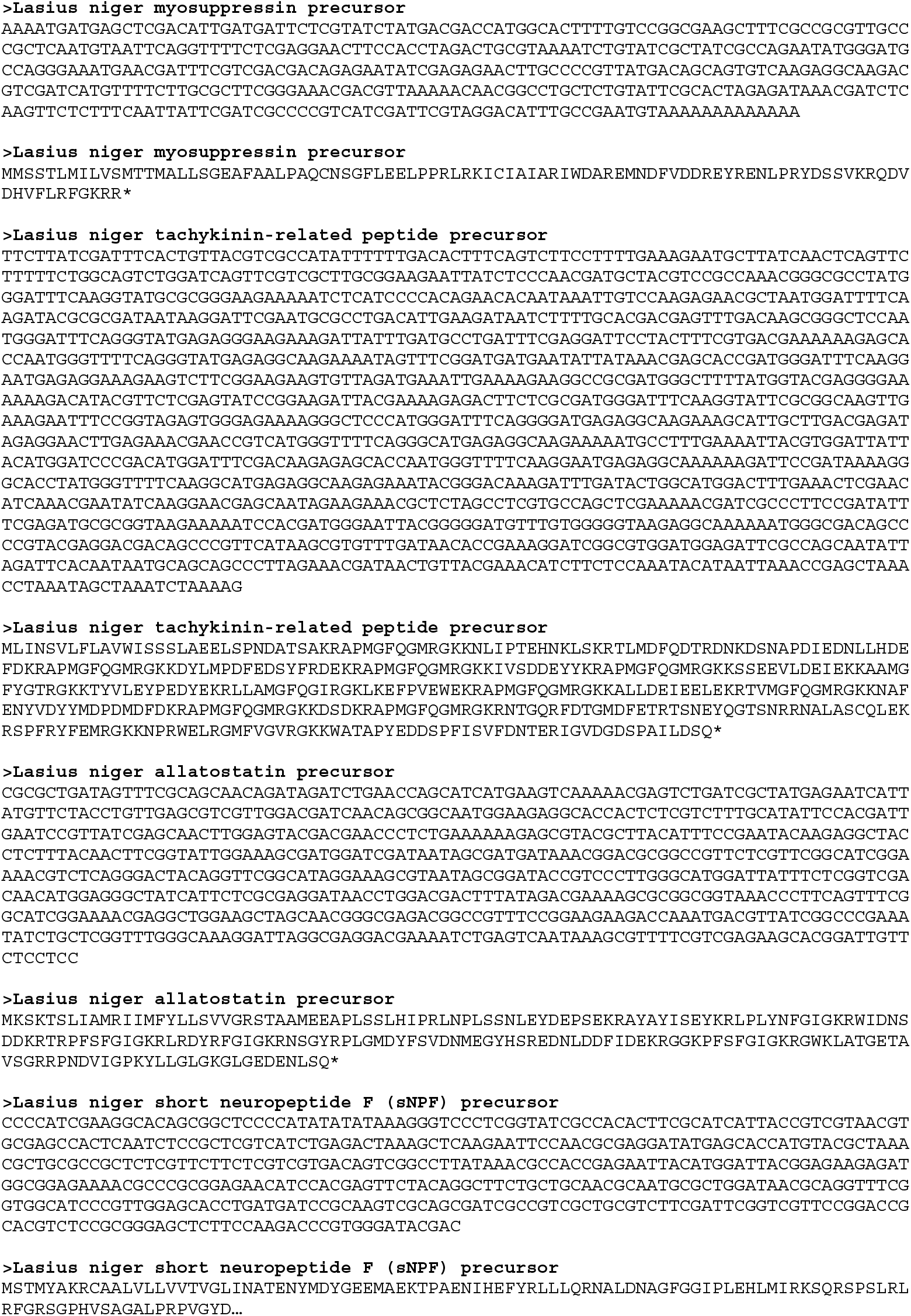
Sequenced full or partial neuropeptides precursors (myosuppressin, tachykinin-related peptide, allatostatin and short neuropeptide F) of *A. sexdens or L. niger*.

